# Golgi fragmentation - One of the earliest organelle phenotypes in Alzheimer’s disease neurons

**DOI:** 10.1101/2022.12.09.519571

**Authors:** Henriette Haukedal, Giulia I. Corsi, Veerendra P. Gadekar, Nadezhda T. Doncheva, Shekhar Kedia, Noortje de Haan, Abinaya Chandrasekaran, Pia Jensen, Pernille Schiønning, Sarah Vallin, Frederik Ravnkilde Marlet, Anna Poon, Carlota Pires, Fawzi Khoder Agha, Hans H. Wandall, Susanna Cirera, Anja Hviid Simonsen, Troels Tolstrup Nielsen, Jørgen Erik Nielsen, Poul Hyttel, Ravi Muddashetty, Blanca I. Aldana, Jan Gorodkin, Deepak Nair, Morten Meyer, Martin Røssel Larsen, Kristine Freude

**Affiliations:** Department of Veterinary and Animal Sciences, Faculty of Health and Medical Sciences, University of Copenhagen, Frederiksberg 1870, Denmark; Center for Non-coding RNA in Technology and Health, University of Copenhagen, 1871 Frederiksberg, Denmark; Novo Nordisk Foundation Center for Protein Research, University of Copenhagen, 2200 Copenhagen, Denmark; Centre for Neuroscience, Indian Institute of Science, Bangalore 560012, India; Copenhagen Center for Glycomics, Department of Cellular and Molecular Medicine, Faculty of Health and Medical Sciences, University of Copenhagen, 2200 Copenhagen, Denmark; Department of Biochemistry and Molecular Biology, University of Southern Denmark, 5230 Odense, Denmark; Department of Drug Design and Pharmacology, Faculty of Health and Medical Sciences, University of Copenhagen, 2100 Copenhagen, Denmark; Danish Dementia Research Centre, Dept. of Neurology, Neuroscience Centre, Copenhagen University Hospital - Rigshospitalet, Copenhagen, Denmark; Institute for Stem Cell Science and Regenerative Medicine, Bangalore 560065, India; Department of Neurobiology Research, Institute of Molecular Medicine, University of Southern Denmark, 5000 Odense, Denmark; Department of Neurology, Odense University Hospital, 5000 Odense, Denmark

**Author notes:** Corresponding author: Kristine Freude.

## Abstract

Alzheimer’s disease (AD) is the most common cause of dementia, with no current cure. Consequently, alternative approaches focusing on early pathological events in specific neuronal populations, besides targeting the well-studied Amyloid beta (Aβ) accumulations and Tau tangles, are needed. In this study, we have investigated disease phenotypes specific to glutamatergic forebrain neurons and mapped the timeline of their occurrence, by implementing familial and sporadic human induced pluripotent stem cell models as well as the 5xFAD mouse model. We recapitulated characteristic late AD disease phenotypes, such as increased Aβ secretion and Tau hyperphosphorylation, as well as previously well documented mitochondrial and synaptic deficits. Intriguingly, we identified Golgi fragmentation as one of the earliest AD phenotypes, indicating potential impairments in protein processing and post-translational modifications. Computational analysis of RNA sequencing data revealed differentially expressed genes involved in glycosylation and glycan patterns, whilst total glycan profiling revealed minor glycosylation differences. This indicates general robustness of glycosylation besides the observed fragmented morphology. Importantly, we identified that genetic variants in Sortilin-related receptor 1 (*SORL1*) associated with AD could aggravate the Golgi fragmentation and subsequent glycosylation changes. In summary, we identified Golgi fragmentation as one of the earliest disease phenotypes in AD neurons in various *in vivo* and *in vitro* complementary disease models, which can be exacerbated via additional risk variants in *SORL1*.

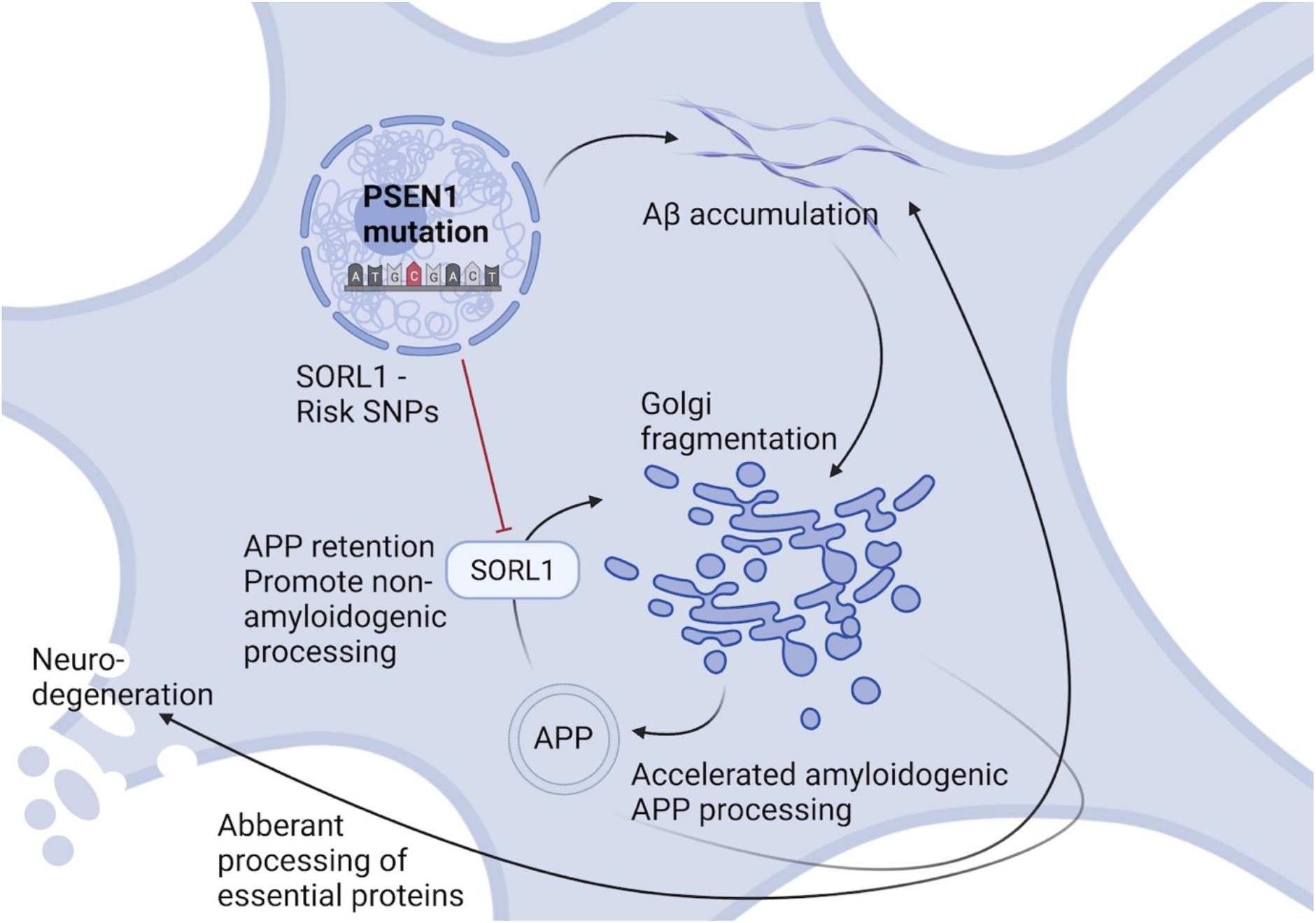

## INTRODUCTION

Alzheimer’s disease (AD) is the most common cause of dementia, accounting for approximately 70% of all cases, with no available curative treatment (1). Familial AD (fAD) presents as early-onset AD and is caused by mutations in *Presenilin 1, Presenilin 2* (*PSEN1, PSEN2*) or *Amyloid Precursor Protein* (*APP*) (2). Although fAD is causative for only 1-5% of AD cases (3), the same pathological hallmarks are shared between fAD and the more abundant multifactorial sporadic AD (sAD).

Most fAD cases carry mutations in *PSEN1*, a catalytic subunit of γ-secretase, responsible for the final cleavage of APP into Amyloid beta (Aβ) peptides. Aβ accumulation leads to aggregation and build-up of extracellular Aβ plaques in patient brains. Together with intracellular neurofibrillary tangles (NFTs), caused by Tau hyperphosphorylation, these are considered classical pathological hallmarks of AD (4–6) and are characteristic in late stages of AD. However, AD pathogenesis initiates decades before these pathologies and symptoms appear, and precise disease mechanisms remain to be elucidated. Mitochondria, metabolic deficits and synaptic dysfunction have been widely studied in relation to AD, and it is evident that all of these are implicated in disease progression (7–10). Nevertheless, the exact timeline and initial triggers of these neuronal phenotypes remain elusive, emphasizing the need to fully understand neuron-specific pathology to remedy the current lack of efficient AD treatments, so far solely focusing on counter-acting Aβ and Tau pathology.

A potential relevant cellular phenotype in AD is Golgi fragmentation, which has recently been reported in various neurodegenerative disorders (11). The Golgi apparatus is the primary site of trafficking, processing, and sorting of most proteins and lipids, potentially linking Golgi fragmentation to abnormal post-translational processing. Alterations in Golgi structure or function can disrupt processing of AD-related molecules, which has been linked to both Aβ and Tau pathologies (12). Moreover, altered trafficking of proteins and metabolites could affect synaptic function and overall neural health. In AD, disruptions of the Golgi stacks have been observed in post-mortem AD brains and transgenic mouse models (13–15), and loss of Golgi ribbons or stack integrity is expected to affect membrane transport, glycosylation and signalling networks (16). These findings led us to investigate if Golgi fragmentation and altered glycosylation act as early events in AD pathogenesis, ultimately promoting neurodegeneration in human induced pluripotent stem cell (hiPSC) derived neuronal fAD models. We further extended our study including sAD hiPSC models, and the *in vivo* 5xFAD mouse model to investigate the universal relevance of this early disease pathology.

## RESULTS

### Generation and Characterization of AD Neurons from Human Induced Pluripotent Stem Cells

Neurons were differentiated from hiPSCs derived from two patients carrying fAD-linked *PSEN1* mutations and their respective isogenic controls corrected via CRISPR/Cas9 gene editing: L150P and L150P GC (gene corrected) (17,18), A79V and A79V GC (19,20), as well as a healthy control-(K3P53) (21) and a CRISPR/Cas9 knock-in hiPSC line carrying the Swedish *APP* mutation (BioSweden) (22). fAD- and control-hiPSCs were successfully differentiated into cortical, glutamatergic forebrain neurons (23) **(Figure 1A)**. Neurite outgrowth analysis revealed a non-significant tendency towards reduced neurite length in the AD lines compared to their respective isogenic controls after one week of differentiation **(Figure 1B, C)**. Neural outgrowth capacity was initially lower for A79V and A79V GC, potentially indicating innate differences between the two hiPSC lines during the early stages of differentiation. However, all hiPSC derived neurons displayed comparable differentiation potential and maturity levels at week seven. This was assessed via expression of microtubule associated protein 2 (MAP2) and Tau **(Figure 1D)**. Astrocyte content in our cultures ranged from 5-10%, detected by Glial Fibrillary Acidic Protein (GFAP) expression **(Figure 1E)**. Most of the cultured neurons were glutamatergic neurons (VGLUT) with few GABAergic neurons (VGAT) **(Figure 1E**, Supplementary, Figure S1).

**Figure 1.**
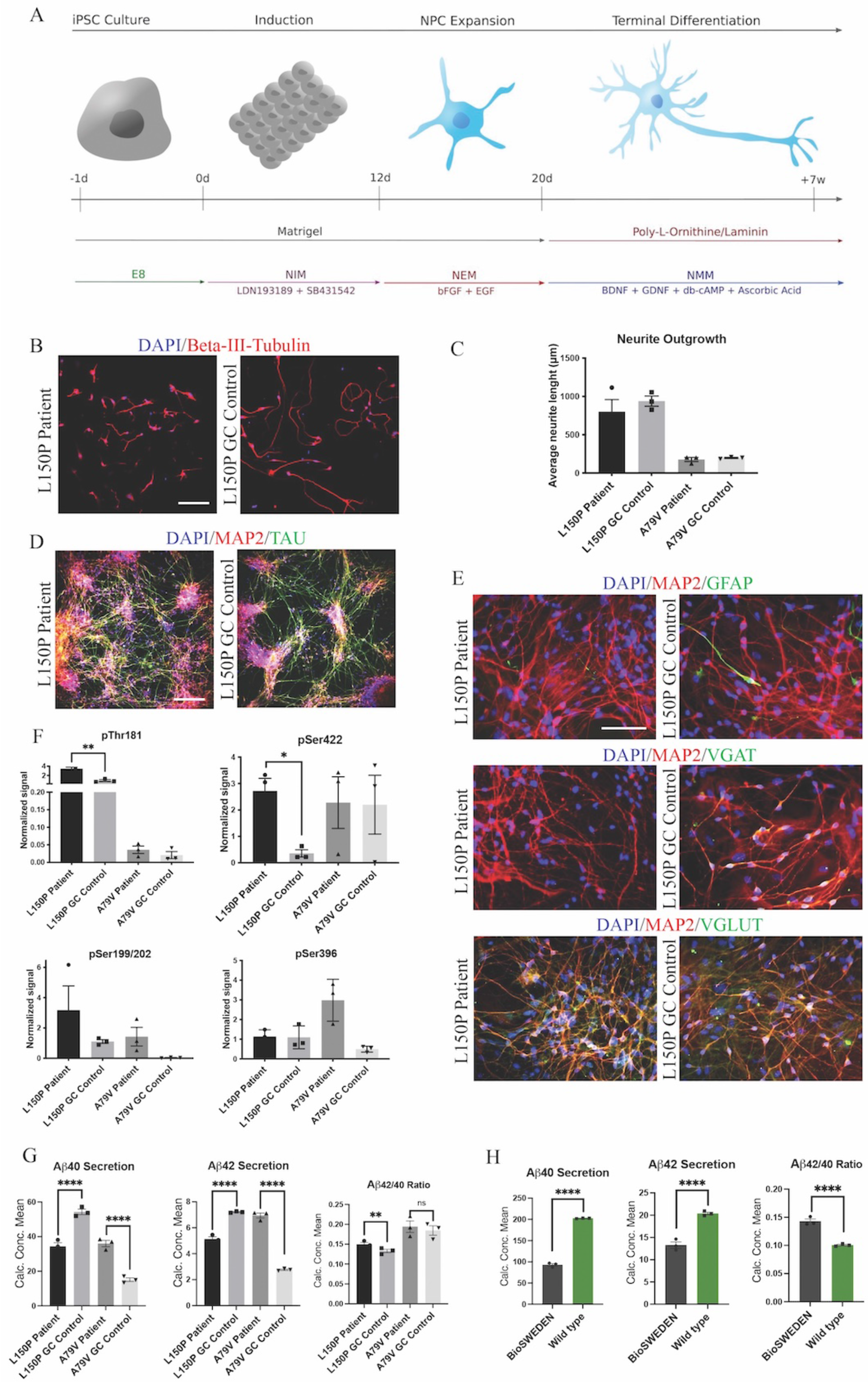
Generation and characterization of hiPSC derived neurons. **A**. Schematic overview of the neural differentiation protocol. **B**. Neurite outgrowth analysis via ICC expression of Beta-III-tubulin. Scale bar 100 µm. **C**. Quantitative assessment of average neurite outgrowth. **D**. Representative ICC images of neuronal markers MAP2 and TAU. Scale bar 100 µm. **E**. Representative ICC images of MAP2, astrocytic marker GFAP, GABAergic neuron marker VGAT and glutamatergic neuron marker VGLUT. Scale bar 50 µm. **F**. Quantitative assessment of Tau phosphorylation (all four isoforms) - pThr181, pSer422, pSer199/202 and pSer396. **G**. Quantitative assessment of Aβ40 and 42 secretion, and Aβ42/40 ratio in L150P and A79V hiPSC derived neurons. **H**. Quantitative assessment Aβ40 and 42 secretion, and Aβ42/40 ratio in K3P53 and BioSweden hiPSC derived neurons. Results are displayed as mean ± standard error of the mean (SEM) from three replicates. Significance levels are indicated by p < 0.05*, p < 0.01**, p < 0.001*** and p < 0.0001****.

### AD Neurons Display Amyloid Beta- and Tau Pathologies

Western blot (WB) analyses revealed significantly increased levels of phosphorylated Tau (p-Tau) isoforms pThr181 and pSer422 in the L150P neurons. A similar trend for pSer199/S202 was observed, but no difference was detected for pSer396. A79V demonstrated similar tendencies but showed no difference in pSer422 levels **(Figure 1F)**. Mesoscale V-PLEX assessment of Aβ peptides 1-40 and 1-42 revealed a significant increase in Aβ42/40 ratio in L150P neurons, likely due to a substantial decrease in Aβ40. This increased ratio was not significant in the A79V neurons; however, up-regulated secretion of both Aβ40 and Aβ42 was detected **(Figure 1G)**, which aligns with other studies in different model systems of the A79V mutation (24–26). Recapitulation of characteristic Aβ and Tau pathologies in our hiPSC derived cell models validates their relevance as *in vitro* AD models. Moreover, increased Aβ42/40 ratio was detected in BioSweden neurons, indicating that other fAD-linked mutations result in similar neuronal phenotypes (**Figure 1H**).

### AD Neurons Display Abnormal Mitochondria Morphology and Distribution

Next, we assessed the mitochondria ultrastructure in fAD and control neurons using transmission electron microscopy (TEM). Here, we demonstrated abnormal cristae-lacking organelle morphology **(Figure 2A)** and a significant increase of relative cristae-less mitochondria to cytoplasm ratio in both L150P and A79V neurons **(Figure 2B)**. The relative individual mitochondria area was unaltered, indicating increased number of cristae-less organelles, and not mitochondria size **(Figure 2C)**. TEM analysis revealed a perinuclear mitochondria accumulation in fAD neurons. MitoTracker™ confirmed a disrupted mitochondria distribution with a significant reduction in mean area of distribution in both L150P and A79V **(Figure 2D, E)**. Rescue of mitochondria phenotypes in our isogenic controls shows that our findings can be attributed to the *PSEN1* mutations. The same distribution pattern was demonstrated in BioSweden neurons, indicating that mitochondria defects are equally relevant for both *PSEN1* and *APP* fAD-linked mutations **(Figure 2F)**. Aberrant mitochondria morphology suggests impaired mitochondria function since the ATP production via electron transport chain takes place within the inner membrane, which depends on an intact membrane and correct cristae folding (27). Abnormalities in energy production and metabolism were verified by computational analysis of RNA sequencing, which revealed differentially expressed genes related to mitochondria and oxidative stress, further validated by qPCR **(Figure 2G, H)**, and supported by proteomics (Supplementary, Table S2C). *CKMT1*, central to metabolism and energy transduction, was significantly increased in both L150P and A79V neurons **(Figure 2G**, Supplementary, Table S2C). Such upregulation is commonly seen in conditions with compromised energy state (28), and might reflect a compensatory mechanism in response to impaired mitochondria function. Moreover, downregulation of *SCARA3*, normally protecting against reactive oxygen species (ROS), was detected in both L150P and A79V neurons (**Figure 2G**, Supplementary, Table S2C), indicating an impaired ability to handle oxidative stress and causing excessive neurotoxic effects (29). Combined, these findings validate mitochondria dysfunction early-on in AD pathogenesis in our seven-week fAD neurons, which are supported by previously described mitochondria phenotypes in the 5xFAD mouse model (Andersen et al., 2021).

**Figure 2.**
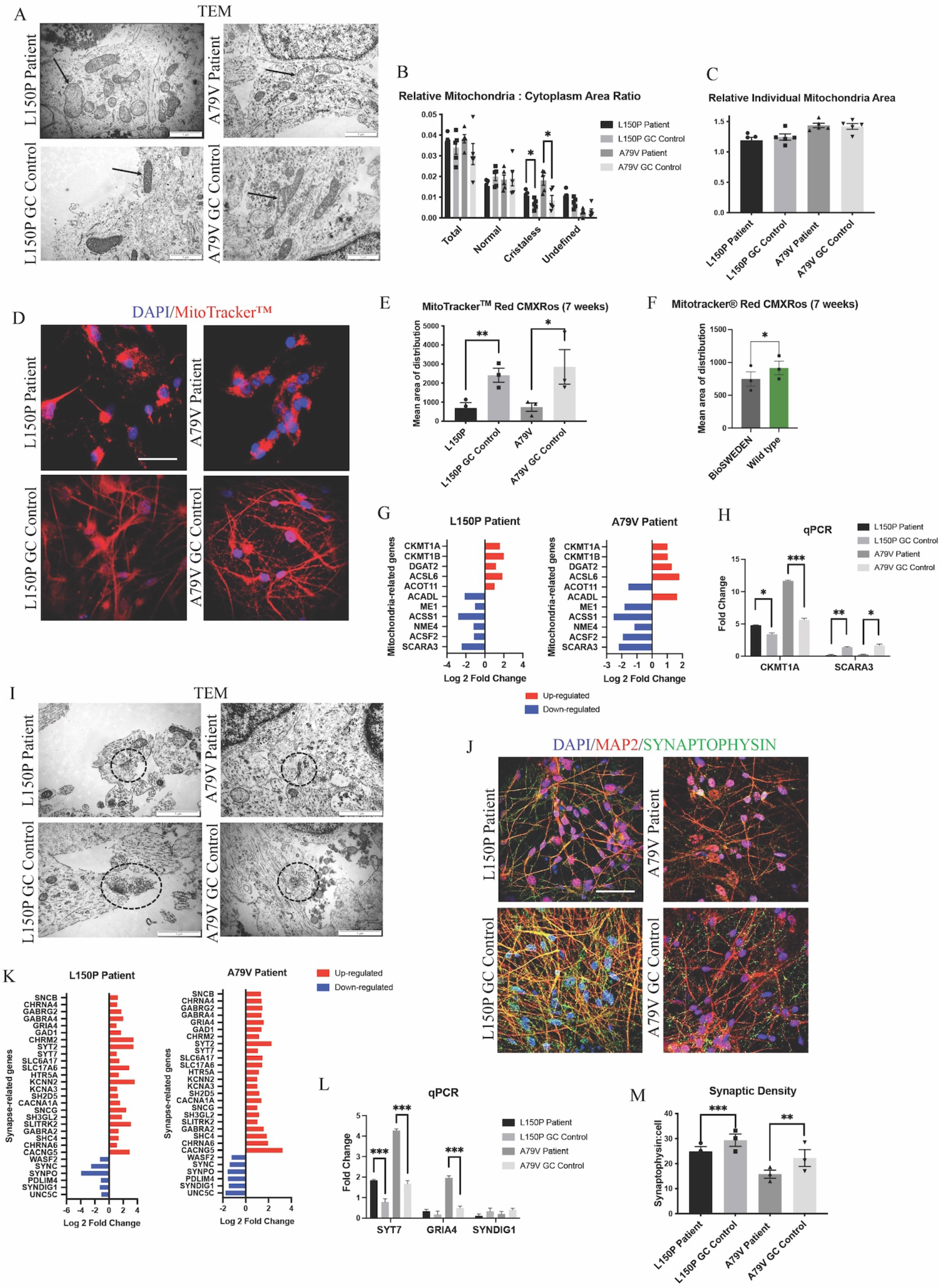
fAD neurons display mitochondria and synaptic deficits. **A**. TEM evaluation of mitochondria (indicated by arrows) ultrastructure. Scale bar 1 µm. **B**. Quantitative TEM morphometry assessment of relative mitochondria to cytoplasm ratio. **C**. Quantitative TEM morphometry evaluation of relative individual mitochondria area. **D**. Representative ICC images of MitoTracker™ analysis. Scale bar 50 µm. **E**. Quantitative assessment of MitoTracker™ analysis showing mean area of distribution in L150P and A79V hiPSC derived neurons. **F**. Quantitative assessment of MitoTracker™ analysis showing mean area of distribution in K3P53 and BioSweden hiPSC derived neurons. **G**. Overview of top significantly differentially expressed genes associated with mitochondria. **H**. qPCR validation of key differentially expressed genes associated with mitochondria identified by RNA-seq analysis. **I**. TEM evaluation of synapses and synaptic vesicles (circled). Scale bar 1 µm. **J**. Representative ICC images of synaptophysin expression. Scale bar 50 µm. **K**. Overview of top significantly differentially expressed genes associated with synapses. Adjusted p-value is displayed in Supplementary, Table S2C. **L**. qPCR validation of key differentially expressed genes associated with synapses identified by RNA-seq analysis. **M**. Quantitative assessment of synaptophysin expression. Results are displayed as mean ± standard error of the mean (SEM) from three replicates. Significance levels are indicated by p < 0.05*, p < 0.01**, p < 0.001*** and p < 0.0001****.

### AD Neurons Display Reduced Synaptic Density

TEM assessment further revealed a reduced number of synapses and synaptic vesicles in L150P and A79V neurons **(Figure 2I)**. Together with reduced synaptophysin expression **(Figure 2J, M)**, these findings indicate reduced synaptic density. Interestingly, transcriptome analysis revealed mainly upregulation of synapse-related genes **(Figure 2K)**, validated by qPCR **(Figure 2L)** and proteomics analysis (Supplementary, Table S2C), suggesting altered synaptic function. Such upregulation has been demonstrated in mild cognitive impairment (MCI) and early AD, potentially acting as a compensatory mechanism for rebalancing synaptic transmission in early-stage AD (31). The fact that our *in vitro* cell models display both mitochondria and synaptic deficits, in addition to Aβ and Tau pathology, further stresses their potential in AD modelling, to better understand early cell-type specific disease perturbations.

### AD Neurons Display Golgi Fragmentation

Since Golgi fragmentation has been identified in neurodegenerative disorders (11), we next investigated Golgi organization and function in L150P and A79V neurons. TEM analysis revealed an abnormal Golgi morphology with dilated cisternae and shortened Golgi stacks **(Figure 3A)**. ICC analysis confirmed a disorganized Golgi pattern in L150P and A79V, compared to a centred Golgi at one pole in the perinuclear region in the isogenic controls, both for the cis-(GM130) (**Figure 3B)**, and trans-Golgi (TGN46) **(Figure 3C)**, indicating a total fragmentation, with an increase in cis- and trans-Golgi surface area **(Figure 3D)**. This was particularly prominent for the trans-Golgi, with significant changes in both mutations at seven weeks. Our findings ware mirrored in BioSweden neurons **(Figure 3E)**, demonstrating Golgi fragmentation as a characteristic phenotype common for both *APP* and *PSEN1* fAD mutations. Moreover, Golgi abnormalities were consistent and present in other model systems. Multicolour Airyscan super-resolution imaging of Neuro 2A (N2A) cells, transfected with human *APP (hAPP)* Swedish, confirmed cis-(GM130) and trans-Golgi (γ-adaptin) fragmentation, with significant increase in average intensity and organelle volume **(Figure 3F, G, H)**. Moreover, Golgi fragmentation was present in two independent male and female sAD hiPSC derived neuronal models **(Figure 3I)**. Likewise, Golgi fragmentation was demonstrated in the 5xFAD transgenic mouse model **(Figure 3J)**. We thus validated our findings in multiple fAD and sAD *in vitro*, as well as fAD *in vivo* models, indicating Golgi fragmentation as a universal early event in AD pathology.

**Figure 3.**
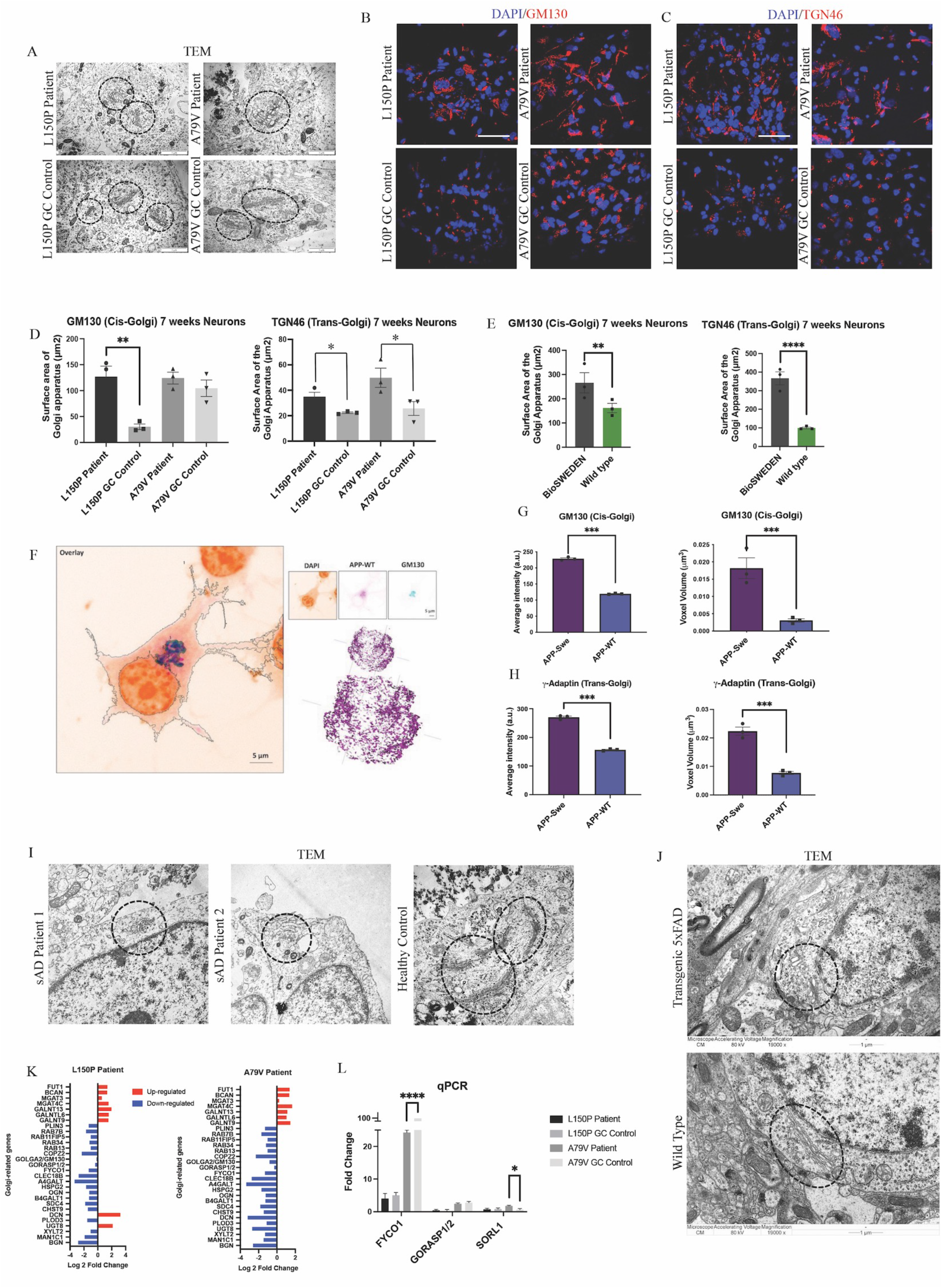
Golgi fragmentation is a universal phenotype in AD. **A**. TEM evaluation of Golgi morphology (circled). Scale bar 1 µm. **B**. Representative ICC images of cis-Golgi marker GM130. Scale bar 50 µm. **C**. Representative ICC images of trans-Golgi marker TGN46. Scale bar 50 µm. **D**. Quantitative assessment of cis- and trans-Golgi surface area in L150P and A79V hiPSC derived neurons. **E**. Quantitative assessment of cis- and trans-Golgi surface area in K3P53 and BioSweden hiPSC derived neurons. **F**. Representative images of cis-Golgi marker GM130 in APP-WT/Swe transfected N2A cells obtained using Airyscan super resolution microscopy. Scale bar 5 µm. **G**. Quantitative assessment of biophysical (intensity) and morphological (voxel volume) traits of cis-Golgi in APP-WT/Swe transfected N2A cells. **H**. Quantification of biophysical (intensity) and morphological (volume) traits of trans-Golgi in APP-WT/Swe transfected N2A cells. **I**. TEM evaluation of dilated Golgi morphology (circled) in hiPSC derived neurons derived from sporadic AD (sAD) patients. Scale bar 1 µm. **J**. TEM evaluation of Golgi fragmentation (circled) in a transgenic 5xFAD mouse model. **K**. Top significantly differentially expressed genes associated with the Golgi apparatus. **L**. qPCR validation of key differentially expressed -Golgi-associated genes identified by RNA-seq analysis. Results are displayed as mean ± standard error of the mean (SEM) from three replicates. Significance levels are indicated by p < 0.05*, p < 0.01**, p < 0.001*** and p < 0.0001****.

Differential expression of Golgi-related genes in L150P and A79V neurons **(Figure 3K)**, validated by qPCR **(Figure 3L)**, further supported Golgi abnormalities. This included *GM130* and *GORASP*, involved in organization, assembly, and vesicle fusion, and *FYCO1*, containing a Golgi dynamics domain facilitating interactions and sorting. Additional differentially expressed genes related to glycosylation, which could indicate impaired Golgi function, include *GALNTs, MAN1C1* and *MGATs* (Supplementary, Table S2C).

### Golgi Fragmentation is an Early Trigger in AD Pathogenesis

To investigate the initial triggers and timeline of AD pathogenesis we assessed our hiPSC derived neurons at both five and seven weeks during the terminal differentiation phase. At week five, the mitochondria distribution remained intact **(Figure 4A)**, whilst Golgi fragmentation was clearly present and particularly prominent in A79V **(Figure 4B)**. Interestingly, the cis-Golgi surface area was significantly increased in five-week A79V neurons, opposed to the seven-week neurons, suggesting a compensatory mechanism that could contribute to maintain Golgi structure and function. Importantly, our results suggest that Golgi fragmentation is one of the earliest neuronal AD events, preceding mitochondria deficits.

**Figure 4.**
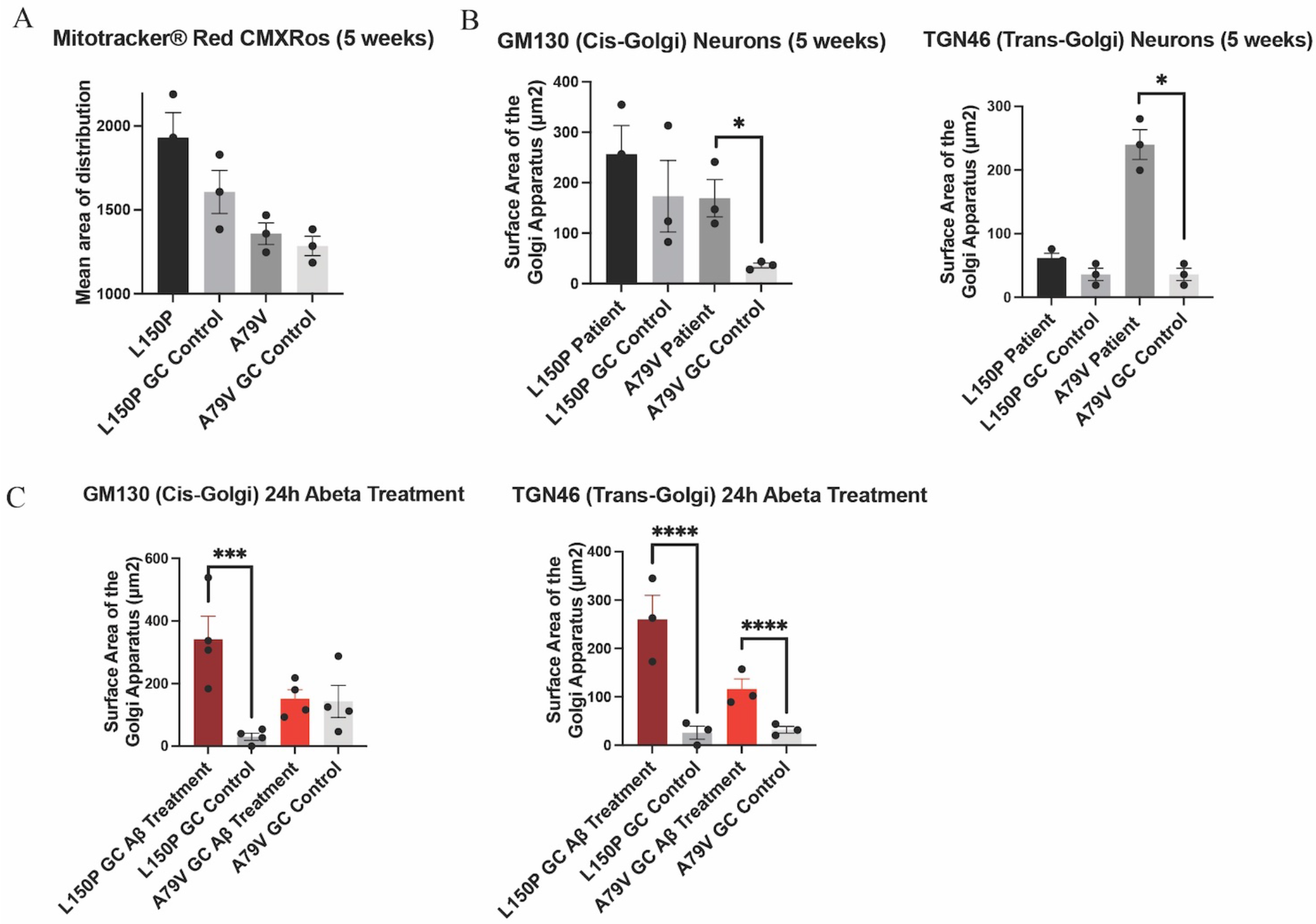
Golgi fragmentation is an early event in AD pathogenesis, triggered by Aβ42 peptides. **A**. Quantitative assessment of MitoTracker™, mean area of distribution, in five-week L150P and A79V hiPSC derived neurons. **B**. Quantitative assessment of cis- and trans-Golgi surface area in five-week L150P and A79V hiPSC derived neurons. **C**. Quantitative assessment of cis- and trans-Golgi surface area in isogenic controls, following a 24-hour Aβ treatment. Results are displayed as mean ± standard error of the mean (SEM) from three replicates. Significance levels are indicated by p < 0.05*, p < 0.01**, p < 0.001*** and p < 0.0001****.

### Golgi Fragmentation can be Induced by Amyloid Beta Peptide Accumulation

Aβ-mediated phosphorylation of the Golgi-matrix protein GRASP65 has been indicated as a trigger of Golgi fragmentation (13). Additionally, Golgi organization is dependent on an intact cytoskeleton, and Tau hyperphosphorylation might exacerbate the disorganization. Fragmentation can lead to a vicious cycle enhancing Aβ and Tau pathologies, ultimately promoting neurodegeneration (12,13). To confirm if increased Aβ is the mechanism underlying the observed Golgi fragmentation, we treated seven-week isogenic control neurons with Aβ42 peptide for 24 hours. Strikingly, this treatment induced fragmentation of both cis-and trans-Golgi, mirroring the seven-week fAD neurons **(Figure 4C)**, clearly indicating that Aβ42 is a trigger of fragmentation. The potential contribution of Tau phosphorylation is highlighted by the fact that L150P neurons demonstrated significant Tau hyperphosphorylation and consequently displayed a more profound fragmentation of the cis-Golgi compartment.

### Golgi Fragmentation is not Directly Affecting Glycosylation Processing

The observed Golgi fragmentation together with changes in genetic expression suggests compromised Golgi function. Therefore, we investigated glycosylation via total glycan profiling of our hiPSC derived neurons. Overall Golgi fragmentation appeared to have little impact, indicating a robust glycosylation machinery, and no consistent changes were observed in *N*-glycan profile for L150P and A79V. However, a few alterations could be observed within the L150P neurons, including a tendency of reduced high-mannose levels, low antenna fucosylation, bisecting GlcNAc and LacdiNAc levels, increased antenna galactosylation **(Figure 5A)**, as well as a sialic acid (SA) linkage shift from 2,3 SA to 2,6 SA **(Figure 5B)**. The *N*-glycan profile of the A79V, as well as the BioSweden (Supplementary, Figure S3), remained unchanged. A similar trend was observed in the *O*-glycan profile. L150P deviated slightly, showing an altered fucosylation pattern, displayed as absent H2N2F1 **(Figure 5C)**, similar to the changes in antenna fucose *N*-glycans. These are likely regulated by the same enzymes, related to terminal glycosylation. Interestingly, L150P and A79V showed a comparable tendency towards increased core 2 *O*-GalNAc glycans **(Figure 5C)**, which could imply that some stages of complex glycosylation could be affected by Golgi fragmentation. Taken together, the glycan alterations observed in the L150P neurons are likely to be disease-associated, but not exclusively linked to *PSEN1* mutations or the Golgi fragmentation.

**Figure 5.**
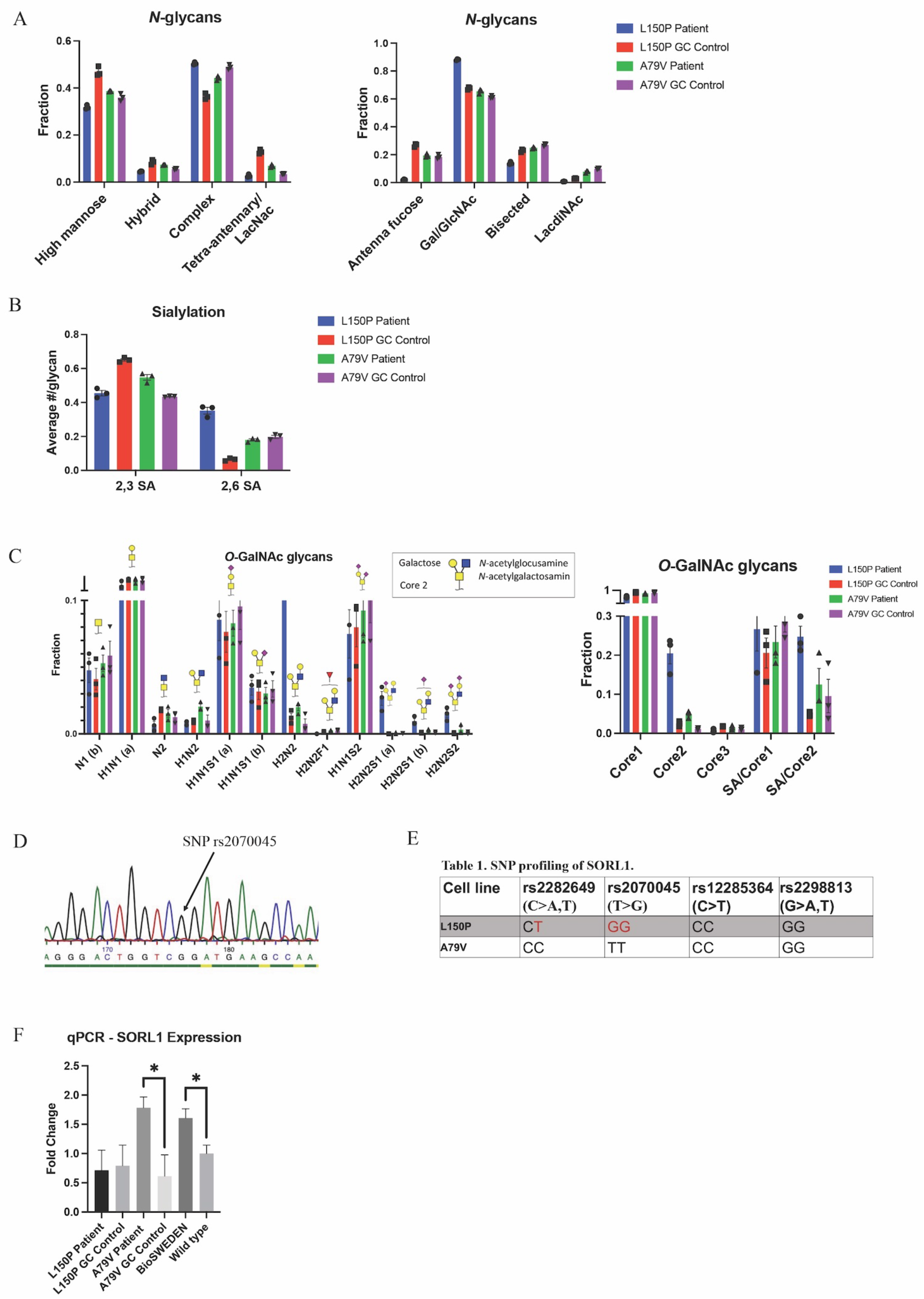
Golgi fragmentation is not directly affecting Golgi function, which is likely linked to multiple mechanisms. **A**. *N*-glycan profiling. **B**. N-glycan ialylation pattern. **C**. *O*-glycan profiling. **D**. Example of SNP profiling via Sanger sequencing. **E**. SNP profiling of the *SORL1* gene. **F**. qPCR validation of *SORL1* expression. Results are displayed as mean ± standard error of the mean (SEM) from three replicates. Significance levels are indicated by p < 0.05*, p < 0.01**, p < 0.001*** and p < 0.0001****.

### Multiple Mechanisms Underlie Golgi Function

The finding of only minor glycan changes in A79V compared to L150P neurons suggests that other underlying mechanisms could maintain the function of the Golgi, despite fragmentation. Sortilin-related receptor 1 (*SORL1*) encodes a protein directly involved in APP processing, and mediates retrograde transport of APP between the trans-Golgi network (TGN) and early endosomes (32). Both genetic and functional alterations of SORL1 has been linked to AD (33,34), and single nucleotide polymorphisms (SNPs) in *SORL1* can confer increased risk of developing AD. Wild-type *SORL1* is suggested to have a protective effect (33,34). Therefore, we assessed the genetic variants of *SORL1* in L150P and A79V hiPSC, via Sanger sequencing **(Figure 5D)** of the genomic locations within the SORL1 protein-coding region, harbouring four SNP sites correlated to AD risk: rs2282649, rs2070045, rs12285364 and rs2298813. Strikingly, we identified the presence of two of these SNP variants in L150P (rs2282649 heterzygote & rs2070045 homozygote), which might contribute to the functional Golgi alteration, causing altered protein trafficking within the trans-Golgi. No *SORL1*-linked SNPs were identified in the A79V **(Figure 5E)**, wild-type K3P53 or knock-in BioSweden hiPSCs. Previous studies have reported decreased *SORL1* expression in AD (35). Instead, A79V and BioSweden neurons showed upregulated *SORL1* expression **(Figure 5F)**. This suggests a beneficial compensatory mechanism that retain Golgi function, which could be characteristic for early stages of AD.

## DISCUSSION

Mutations in *PSEN1* are the most common causes of fAD, but their precise function has yet to be fully elucidated. Here, we investigated phenotypes related to two fAD linked *PSEN1* mutations, as well as the *APP* Swedish mutation, in neurons derived from hiPSC. These models recapitulate characteristic disease hallmarks such as Aβ, Tau, mitochondria and synaptic pathology (36,37). More recently, Golgi fragmentation has been identified in AD models, including post-mortem AD brains (12,15). To date, it was not clear if mitochondria dysfunction precedes Golgi fragmentation or vice versa. Here, we report that Golgi fragmentation is evident prior to mitochondria deficits in our models, placing it as one of the initial events in AD neural pathology. Importantly, we show that fragmentation is consistent in both fAD and sAD, and throughout multiple *in vitro* and *in vivo* model systems, independent of the underlying mutations, reinforcing the relevance of our findings for AD in general. We further indicate that Aβ42 peptides can act as a triggering mechanism, suggesting a timeline where Aβ accumulation is sufficient to induce Golgi fragmentation, followed by mitochondria and synaptic dysfunction.

Glycosylation, which largely takes place in the Golgi, is the most common form of post-translational modification, and glycan changes have been identified in AD (38). To evaluate if Golgi function was impaired in a broader sense, we investigated the glycan pattern in our fAD neurons. Surprisingly, Golgi fragmentation had little impact on glycosylation, indicating a certain robustness of the glycosylation system. Remarkably, some changes in glycan patterns were evident for the L150P neurons, both within the *N*- and *O*-glycan profiles, revealing that the functional outcome of Golgi fragmentation is dependent on the underlying genetic profile of affected individuals and not only on mutations in *PSEN1* or *APP*. The glycan alterations found in L150P neurons mainly affected the final decoration of the sugar structures, which largely takes place in the trans-Golgi. There was no significant concordance between the glycogene candidates identified via RNA sequencing analysis and the global neuron glycosylation patterns. This suggests that the glycosylation in this study is not regulated at the level of the glycogene expression but rather on enzyme localization and carrier glycoprotein production. Such discrepancies between RNA networks and proteome are not uncommon and have previously been described in AD brains (39). Importantly, fragmentation was more prominent in the L150P neurons, whilst the cis-Golgi was not significantly affected in A79V neurons, potentially reflecting the intact *N-*glycan profile. Reduced levels of high-mannose in L150P neurons supports this theory, as mannosidase activity occurs in the cis-Golgi (40). Golgi fragmentation has been suggested to accelerate protein trafficking through increased surface area for vesicle budding and altered protein sorting (13). Cis-Golgi fragmentation could hence contribute to increased mannosidase activity, explaining the reduced high-mannose levels only evident in L150P. Moreover, the *O*-glycan pattern in L150P neurons, which deviates slightly from the other lines, might explain the significant changes we observe in its p-Tau levels. Hence, the mainly intact *O*-glycosylation pattern in the A79V neurons could result in non-significant changes in Tau phosphorylation.

The only consistent trend in glycan pattern was upregulation of the complex *O*-glycan structure core 2, one of the processes that mainly occurs in the trans-Golgi. The mutually significant fragmentation of this compartment could thus underlie these effects. As hypothesized, if fragmentation enhances protein processing through increased vesicle budding and impaired protein sorting, it could alter the localization of proteins and their respective processing enzymes and fail to sort them into separate compartments, thus accelerating the post-translational processing, ultimately causing altered glycan patterns. However, the findings of a robust glycosylation process in A79V, BioSweden as well as Aβ-treated control neurons, suggest that although Aβ is sufficient to trigger Golgi fragmentation, multiple mechanisms might underlie Golgi function, and prolonged fragmentation might be needed to ultimately induce a functional effect. Moreover, although Golgi fragmentation had no direct or exclusive impact on overall glycosylation, other post-translational modification processes could potentially be affected.

Golgi fragmentation was dependent on the individual genetic profile, suggesting involvement of multiple mechanisms. SORL1, which is both genetically and functionally linked to AD risk, retains APP in the TGN, and can potentially exert a protective effect against Aβ toxicity (32). SORL1 and APP co-localize in the Golgi and endosomal compartments. Neural overexpression of SORL1 leads to redistribution of APP to the Golgi and reduced amyloidogenic processing, whereas SORL1 depletion causes increased Aβ production (Andersen et al., 2005). The extracellular plasma membrane has been considered the major site for α-secretase activity. However, recent studies in primary neurons have demonstrated α-secretase activity in the TGN, with increased levels of C83 (α-CTF) compared to C99 (β-CTF) following APP accumulation (42). This indicates TGN as a major additional site for α-secretase processing. SORL1-mediated APP retention in the TGN could thus be a protective mechanism. Mutant SORL1 maintain APP-binding activity, but in contrary to wild-type SORL1 leads to misdirected APP trafficking into non-Golgi compartments that in turn increases Aβ production (43).

We identified several risk-associated SNPs in *SORL1* in our L150P hiPSC, which could contribute to the observed altered Golgi function, absent in the other lines. Interestingly, SORL1 expression was upregulated in both A79V and BioSweden neurons, in contrary to the decreased levels commonly seen in AD patients. These findings suggest a possible compensatory mechanism that might influence the glycosylation process and could potentially be correlated to the differences in Aβ peptides secretion. The Aβ42/40 ratio, which was increased in L150P, is directly correlated to Tau pathology (44). On the contrary, A79V neurons displayed increased levels of individual Aβ40 and 42 levels, possibly linked to the APOE ε4/4 profile of this patient, suggested to increase Aβ production (45). Our observations further indicate a compensatory mechanism of SORL1 that counteracts the increase in Aβ levels early-on in A79V neurons, eventually lowering the impact on perturbations of the Golgi. This is in line with the fragmentation pattern observed for the cis-Golgi, which is significant in five-week A79V neurons but restored at seven weeks.

APP and SORL1 contain multiple glycosylation sites and undergo processing within the Golgi. SORL1-mediated APP retention in the TGN potentially allows for proper processing, despite the compromised structure. Changes in glycosylation pattern of key AD-related molecules such as APP, Tau, BACE1 and APOE have previously been identified (46–50). In this study we focused on global glycan changes, and the analyses account for the total abundance of neuronal glycan patterns and not individual proteins or the cellular localization. While powerful in giving an overall overview, this approach might miss more subtle changes. The global glycosylation effects observed could be driven by altered abundance of specific carrier glycoproteins. Alternatively, multiple proteins could be affected, but the functional relevance of these changes might differ for individual proteins, such as APP and SORL1.

SORL1 has been shown to interact with APOE, especially the ε4 variant. Reduced levels of SORL1 have been observed in neural stem cells from APOE ε4/4 carriers (51), whereas increased levels have been detected in cerebrospinal fluid (CSF) of AD patients (52). A79V neurons carry APOE ε4/4 and BioSweden carry APOE ε3/4. Based on our findings we propose that the presence of APOE ε4 correlates with upregulation of SORL1 expression as a compensatory mechanism. Hence, we observed the opposite in L150P neurons carrying APOE ε2/3, displaying no upregulation of SORL1 expression. SORL1 overexpression has been shown to increase Aβ uptake in an APOE isoform-dependent manner, and more efficiently in the presence of the APOE ε4 isoform (53). Dysfunctional SORL1 (caused by SNP variants) can lead to impaired binding to APP and Aβ. This could in turn alter APP trafficking, reduce SORL1-mediated retention in the TGN, and thus increase Aβ production in our L150P neurons. Moreover, SORL1 has been proposed to influence Tau pathology, and SORL1 SNPs have been linked to increased p-Tau levels (54). This aligns with the significant changes in p-Tau in L150P neurons. It is thus evident that the genetic background of our AD patients impacts their neuronal phenotypes, and other AD-related SNPs might contribute to differences in cellular phenotypes, providing additional AD risk or protective factors besides the presence of *PSEN1* mutations.

Notably, the two patients in the study presented with clinical differences (**Figure 6**). The L150P carrier was a 58-year-old male, with established dementia at the time of sampling and generalized cerebral atrophy, identified through magnetic resonance imaging (MRI). The A79V carrier was a 48-year-old female displaying only slight personality changes, with dementia evolving eight years after sampling. An MRI was not performed for this patient. However, a fluorodeoxyglucose (FDG)-positron emission tomography (PET) scan was normal, whilst a Pittsburgh compound B-PET scan revealed increased cortical binding of amyloid in a pattern typical for fAD-linked *PSEN1* mutations. Pathological levels of Aβ and Tau were detected in her CSF.

**Figure 6.**
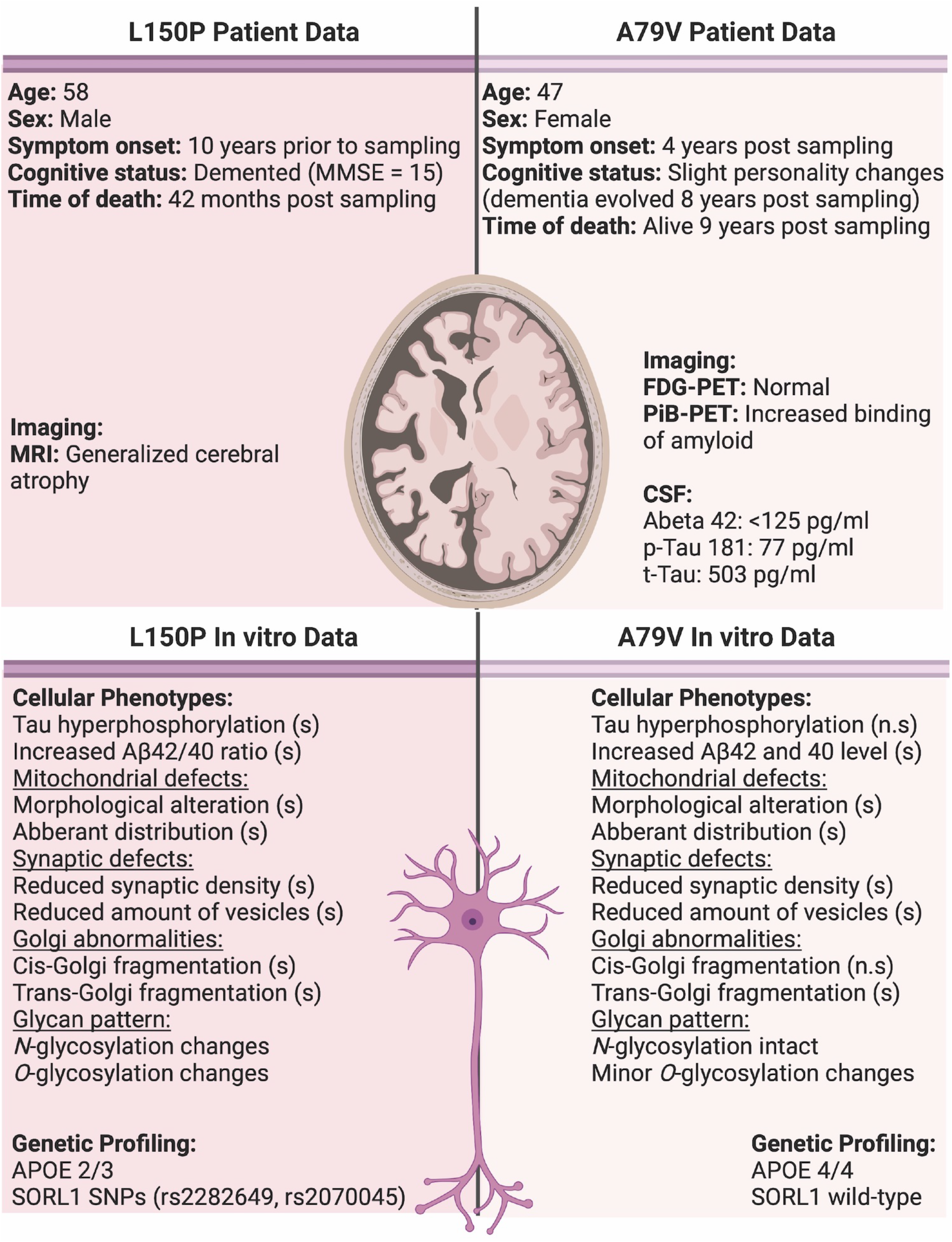
Overview of patient data compared to observations in our *in vitro* cell models. Significant changes are indicated by (s), and non-significant by (n.s). CSF normal reference values; Abeta 42 > 400 pg/ml, p-Tau 181 < 80 pg/ml and t-Tau < 300 pg/ml.

PSEN1 has been suggested to not only act as the catalytic subunit of γ-secretase, but also as a regulator of APP trafficking to the cell surface. fAD-linked mutations have been demonstrated to reduce this APP cell surface delivery, whereas loss-of-function mutations lead to APP accumulation at the plasma membrane (55). *PSEN1* mutations could thus modulate APP processing through multiple mechanisms. The different physical location of the A79V and L150P mutations within *PSEN1* could further contribute to the observed diverse phenotypic outcome. L150P is located closer to the catalytic protein domain, which could exacerbate the neuronal disease phenotype, whilst the A79V mutation is located at the intracellular C-terminal end.

In summary, our findings demonstrate that fAD hiPSC derived neurons display hallmark AD pathologies. We identify Golgi fragmentation as one of the earliest neuronal events in AD pathogenesis and provide insight of a causative mechanism of Aβ accumulation potentially in combination with Tau hyperphosphorylation. We show that glycosylation depends on multiple mechanisms and suggest that the *SORL1* genetic background and expression level could contribute to maintain Golgi functionality. The demonstration of Golgi-related phenotypes in additional model systems clearly indicates it as a universal perturbation in both familial early-and sporadic late-onset AD, validating the clinical relevance of our findings. Pharmaceutical intervention strategies targeting these early processes could be a potential new treatment avenue. Notably, long-term changes in the glycosylation profile could alter the overall cell surface composition, potentially inducing an inflammatory response. A remarkable future aspect of this study would therefore be evaluation of neuron-to-microglia interactions in co-culture systems.

## MATERIALS AND METHODS

For detailed description of experimental procedures see supplementary information.

### hiPSC Generation, Cell Culture and Neural Differentiation

The fAD hiPSC cell lines used in the study (Supplementary, Table S1A) have previously been published (17–22). Generated hiPSC sAD lines (not published) have been characterized and assessed for pluripotency markers and genomic integrity (Supplementary, Figure S2). Neural differentiation was performed according to a modified protocol (23).

### Immunocytochemistry, MitoTracker™, Western Blot, Transmission Electron Microscopy and qPCR

ICC, MitoTracker™, WB, TEM and qPCR were performed according to previously described protocols (56). Image processing and analysis was performed in Fiji ImageJ 2.0.0-rc-65/1.51s.

### Mesoscale Assessment of Aβ Peptide Secretion and Aβ Treatment

Secretion of Amyloid beta (Aβ) peptides; Aβ38, Aβ40 and Aβ42, was assessed using the V-PLEX Plus Aβ Peptide Panel 1 Kit (MSD, 6E10), and the MESO QUICKPLEX SQ 120 Imager (MSD), connected to the DISCOVERY WORKBENCH 4.0 software, according to the manufacturer’s protocol.

### Amyloid Beta Treatment

Seven-week isogenic control- and healthy control hiPSC derived neurons were treated with 2.5 μM Aβ peptide 1-42 (Sigma, PP69-0.25MG) for 24 hours.

### RNA Extraction, bulk RNA Sequencing and Computational Analysis

RNA was extracted from seven-week *PSEN1* fAD- and isogenic control hiPSC derived neurons using the RNeasy® Plus Mini Kit (Qiagen, 74134), according to the manufacturer’s protocol. Library preparation and sequencing (DNBseq, 2×100 nt, stranded paired-end) was done by an external provider (Beijing Genomics Institute, BGI). The computational analysis of the RNA sequencing data is described in detail in a separate study (Corsi, Gadekar et al, in prep.) and in the supplementary information.

### Proteomic Assessment and *N*- and *O*-glycan profiling

Proteomic assessment of seven-week *PSEN1* fAD- and isogenic control hiPSC derived neurons was performed by mass spectrometry, according to a previously published protocol (57). Moreover, for glycan profiling, total cell lysate protein *N*- and *O*-glycans were released and analyzed by C18 nanoflow liquid chromatography (LC) coupled to mass spectrometry (MS) as described previously, with minor adaptations (58), from *PSEN1* fAD-, *APP* fAD-, isogenic control-, control- and Aβ-treated control hiPSC derived neurons.

### N2A cells and 5xFAD Transgenic Mice

Neuro 2A (N2A) cells were transiently transfected with either *APP* Swedish- or wild type h*APP*, using Lipofectamine 2000 (Thermo Fisher Scientific, 11668030) according to a previously described protocol (59). Following, ICC and Airyscan super-resolution microscopy was performed as described previously (60). For further validation, cortex and hippocampus from male 5xFAD transgenic mice (N=5) and male wild-type mice (N=4) were evaluated by TEM analysis.

### Statistical Analysis

Statistical analyses were performed using GraphPad Prism (Version 9.2.0) with default options, and statistical significance was determined using a Student’s *t* test, multiple *t* test or two-way Anova with correction for multiple comparison. Data is presented as mean ± standard error of the mean (SEM) for all experiments with statistical significance *p < 0.05, **p < 0.01, ***p < 0.001 and ****p < 0.0001. Gene differential expression was performed using DESeq2 (Version 1.22.2) (61) using the Benjamini-Hochberg method to adjust Wald test p-values in multiple testing.

## Acknowledgements

The authors would like to thank Nadine Becker-von Buch and Maria Pihl for excellent technical assistance.

## Author contributions

H.H. and K.K.F. performed the experimental design. H.H. performed most of the experiments. S.V., P.S. and F.R.M. performed western blot analysis, M.M. performed the mesoscale analysis, P.J. and M.R.L. performed the proteomics analysis. N.H., F.K.A. and H.W. performed the glycan profiling. G.I.C., N.T.D., V.P.G., and J.G. performed the computational analyses of the RNA sequencing data. S.K and D.N were involved in assessment and airyscan microscopy of N2A cells. S.C. were involved in the supervision of the work and contributed with the qPCR. A.H.S., T.T.N. and J.E.N. provided regulatory approvals, patients’ consent, and clinical data. A.P. and C.P. generated cell lines. A.C., P.H., B.A. and R.M. contributed to the manuscript preparation. All the authors read and approved the final manuscript.

## Ethical statement

This study has been approved by the Ethics Committee of the Capital Region of Denmark (H-4-2011-157). Before enrolment, written informed consent was obtain from all subjects who provided samples for hiPSC generation.

## Data Availability

The datasets supporting the conclusions of this article are included within the article and the additional supplementary source data file. The dataset generated for the RNA sequencing analysis has been deposited in the NCBI GEO database with the accession number GEO: GSE211993.

