## Supplemental files for "Golgi fragmentation - One of the earliest organelle phenotypes in Alzheimer’s disease neurons"

### Table of Contents

**Figure S1.** Generation and characterization of A79V hiPSC derived neurons.

**Figure S2.** Generation, characterization and quality assessment of established sporadic AD hiPSC lines.

**Figure S3.** *N*- and *O*-glycan profiling of wild-type (K3P53), knock-in BioSweden and Aβ-treated wild-type hiPSC derived neurons.

**Table S1A.** Summary of cell lines used in the study.

**Table S1B.** List of antibodies used for immunocytochemistry.

**Table S1C.** List of antibodies used for western blot.

**Table S1D.** List of selected differentially expressed genes identified by bioinformatic analyses of RNA sequencing.

**Table S1E.** List of primers used for qPCR.

**Supplementary Experimental Procedures**

**Supplementary References**

**Supplementary Information – Source Data**

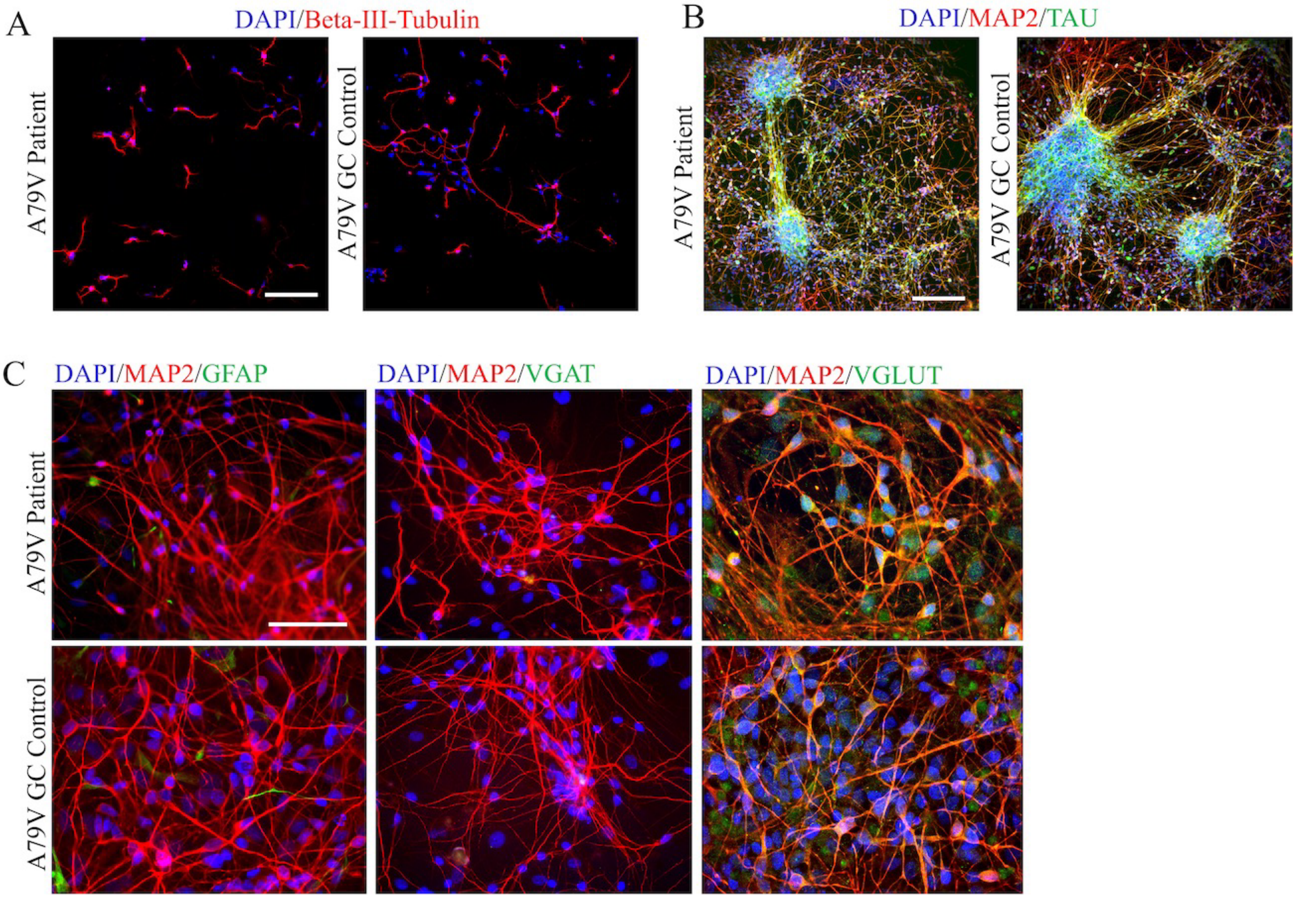


**Figure S1.** Generation and characterization of A79V hiPSC derived neurons. **A.** Neurite outgrowth analysis via ICC expression of Beta-III-tubulin. Scale bar 100 µm. **B.** Representative ICC images of neuronal markers MAP2 and TAU. Scale bar 100 µm. **C.** Representative ICC images of MAP2, astrocytic marker GFAP, GABAergic neuron marker VGAT and glutamatergic neuron marker VGLUT. Scale bar 50 µm.

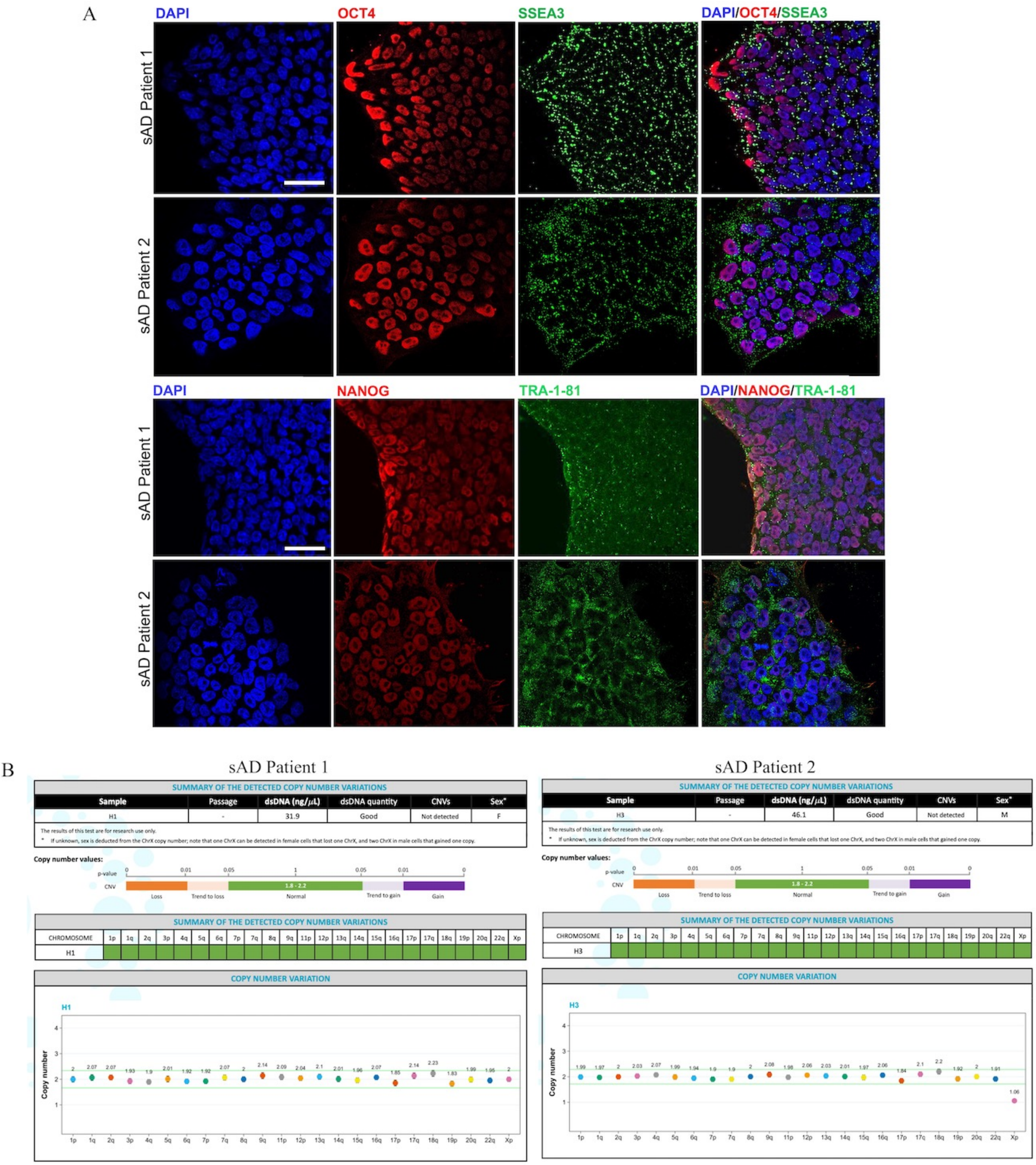


**Figure S2.** Generation, characterization and quality assessment of established sporadic AD hiPSC lines. **A.** Representative ICC images of pluripotency markers OCT4 and NANOG (red), as well as SSEA3 and TRA-1-81 (green). Scale bar 50 µm. **B.** Copy number variance (CNV) analysis.

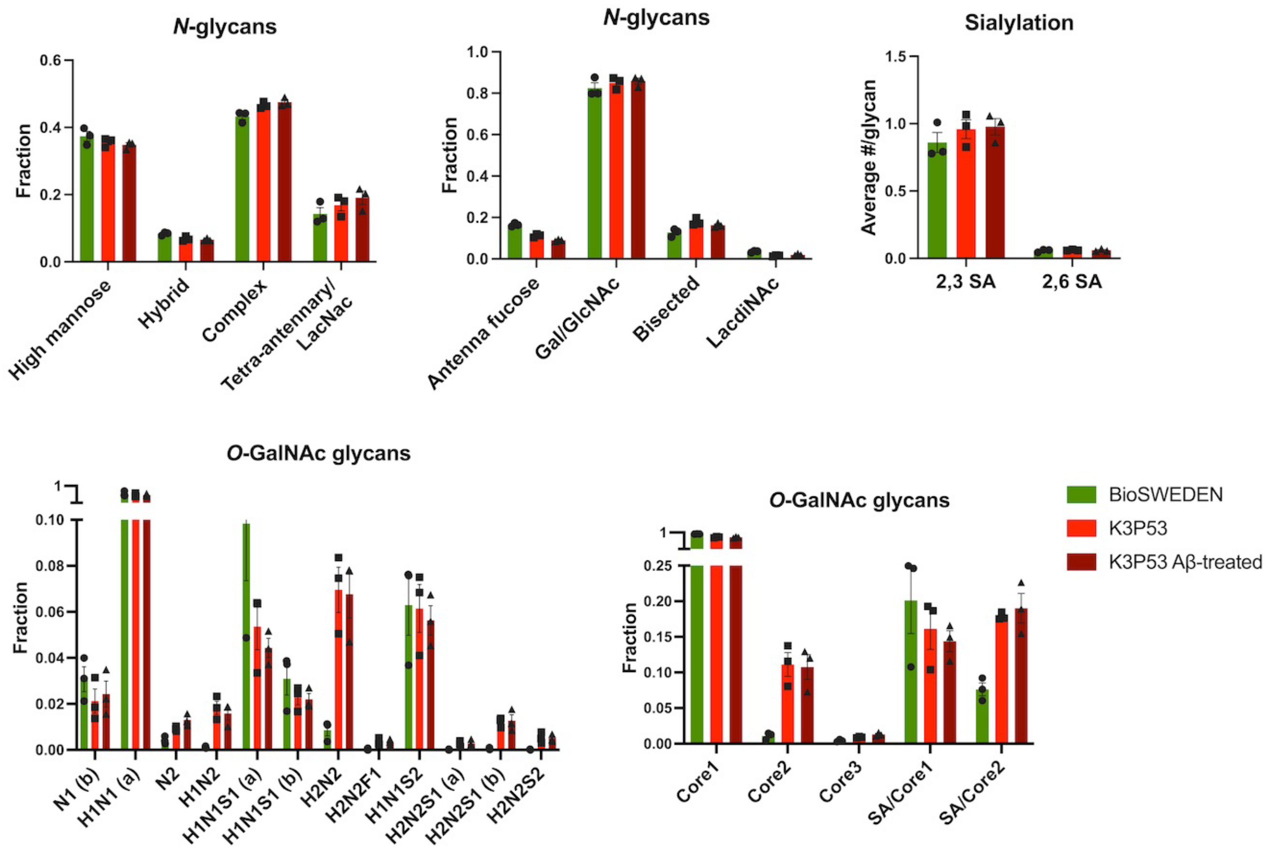


**Figure S3.** *N*- and *O*-glycan profiling of wild-type (K3P53), knock-in BioSweden and Aβ-treated wild-type hiPSC derived neurons.

| hiPSC line | Genotype | Status | Sex | Age | Karyotype | Pluripotency associated marker | Reference |
|---|---|---|---|---|---|---|---|
| L150P | c.449 T>C | Symptomatic AD | Male | 58 | Normal | + | (1) |
| L150P GC | Gene correction | Isogenic control | Male | 58 | Normal | + | (2) |
| A79V | c.236 C>T | Pre-symptomatic AD | Female | 48 | Normal | + | (3) |
| A79V GC | Gene correction | Isogenic control | Female | 48 | Normal | + | (4) |
| K3P53 | Wild-type | Control | Male | 18 | Normal | + | (5) |

| | | | | | | | |
|---|---|---|---|---|---|---|---|
| Bio SWEDEN | APPswe | Knock-in fAD | Male | 18 | Normal | + | (6) |
| sAD Patient 1 | - | sAD | Female | 71 | Normal | + | n/a |
| sAD Patient 2 | - | sAD | Male | 70 | Normal | + | n/a |

**Table S1A.** Summary of cell lines used in the study. Information on cell lines included in the study, displaying name, genotype, disease status, sex, age, karyotype, expression of pluripotency markers and publication reference.

| Assessment | Antibody | Dilution | RRID | Company and Cat. No. |
|---|---|---|---|---|
| iPSC verification | Goat anti-OCT4 | 1:500 | AB_653551 | Santa Cruz: Sc-8628 |
| | Rabbit anti-NANOG | 1:500 | AB_1268451 | PeproTech: 500-P236 |
| | Rat anti-SSEA3 | 1:100 | AB_1236554 | Biolegend: 330302 |
| | Mouse anti-SSEA4 | 1:100 | AB_1089208 | Biolegend: 330402 |
| NPC verification | Mouse anti-NESTIN | 1:500 | AB_2251134 | Millipore: MAB5326 |
| | Rabbit anti-PAX6 | 1:100 | AB_291612 | Covance: PRB-278P |
| Neurite outgrowth | Mouse anti-TUJ1 | 1:100 | AB_2210524 | Millipore: MAB1637 |
| Neuron verification | Mouse anti-MAP2 | 1:500 | AB_477171 | Sigma: M1406 |
| | Rabbit anti-MAP2 | 1:500 | AB_10807820 | Millipore: AB2290 |
| | Rabbit anti-TAU | 1:200 | AB_10013724 | Dako: A0024 |
| | Rabbit anti-VGLUT | 1:500 | AB_887875 | Synaptic systems: 135303 |
| | Rabbit anti-VGAT | 1:500 | AB_2301998 | Millipore: AB5062P |
| | Rabbit anti-GFAP | 1:500 | AB_2109645 | Millipore: AB5804 |
| Golgi fragmentation | Sheep anti-TGN46 | 1:500 | AB_324049 | Bio-Rad: AHP500 |
| | Mouse anti-GM130 | 1:500 | AB_398141 | BD Biosciences: 610822 |
| | Mouse anti-γ-adaptin | 1:500 | AB_397768 | BD Biosciences: 610385 |
| Synaptic density | Rabbit anti-SY38 | 1:200 | AB_2198854 | Abcam: AB8049 |

| | | | | |
|---|---|---|---|---|
| Secondary antibodies | Alexa Fluor 488 donkey anti-mouse IgG | 1:500 | AB_141607 | Thermo Fisher Scientific: A21202 |
| | Alexa Fluor 488 donkey anti-rabbit IgG | 1:500 | AB_141708 | Thermo Fisher Scientific: A21206 |
| | Alexa Fluor 594 donkey anti-mouse IgG | 1:500 | AB_141633 | Thermo Fisher Scientific: A21203 |
| | Alexa Fluor 594 donkey anti-rabbit IgG | 1:500 | AB_141637 | Thermo Fisher Scientific: A21207 |
| | Alexa Fluor 594 donkey anti-rat IgG | 1:500 | AB_2535795 | Thermo Fisher Scientific: A21209 |
| | Alexa Fluor 594 donkey anti-goat IgG | 1:500 | AB_2534105 | Thermo Fisher Scientific: A11058 |
| | Alexa Fluor 594 donkey anti-sheep IgG | 1:500 | AB_2534083 | Thermo Fisher Scientific: A11016 |

**Table S1B.** List of antibodies used for immunocytochemistry. Overview of antibodies used for the different immunocytochemistry assessments, including both primary and secondary antibodies, as well as information on dilution factors, company, cat. no. and RRID.

| **Assessment** | **Antibody** | **Dilution** | **RRID** | **Company and Cat. No.** |
|---|---|---|---|---|
| Tau phosphorylation | Rabbit anti-phospho-tau Ser199/Ser202 | 1:1000 | AB_2533749 | Thermo Fisher Scientific: 44768G |
| | Mouse anti-PHF-Tau 181 | 1:500 | AB_223651 | Thermo Fisher Scientific: MN1050 |
| | Rabbit anti-tau phospho S422 | 1:500 | AB_1603345 | Abcam: Ab79415 |
| | Rabbit anti-phospho-tau Ser396 | 1:1000 | AB_2533745 | Thermo Fisher Scientific: 44752G |
| Normalization | Rabbit anti-GAPDH | 1:5000 | AB_10167668 | Santa Cruz: sc-25778 |
| | Mouse anti-β-actin | 1:5000 | AB_476744 | Sigma: A5441 |

| | | | | |
|---|---|---|---|---|
| Secondary antibodies | IRDye 800CW donkey anti-mouse IgG | 1:15000 | AB_621847 | LI-COR: 926-32212 |
| | IRDye 680LT donkey anti-rabbit | 1:15000 | AB_10706167 | LI-COR: 926-68023 |

**Table S1C.** List of antibodies used for western blot. Overview of antibodies used for the Tau western blot assessment, including both primary and secondary antibodies, as well as information on dilution factors, company, cat. no. and RRID.

| Compartment/Pathway | Gene | Ensemble ID |
|---|---|---|
| Mitochondria - oxidative stress | SCARA3 | ENSG00000168077 |
| Mitochondria - metabolism | ACSF2 | ENSG00000167107 |
| | NME4 | ENSG00000103202 |
| | ACSS1 | ENSG00000154930 |
| | ACSL6 | ENSG00000164398 |
| | DGAT2 | ENSG00000062282 |
| | ME1 | ENSG00000065833 |
| | ACADL | ENSG00000115361 |
| | ACOT11 | ENSG00000162390 |
| Mitochondria - ATP/energy production | CKMT1B | ENSG00000237289 |
| | CKMT1A | ENSG00000223572 |
| Synapses - receptors | SNAP25 | ENSG00000132639 |
| | GRM7 | ENSG00000196277 |
| | GRIA1 | ENSG00000155511 |
| | SYAP1 | ENSG00000169895 |
| | SYP | ENSG00000102003 |
| | GRIN1 | ENSG00000176884 |
| | CACNG5 | ENSG00000075429 |
| | CHRNA6 | ENSG00000147434 |
| | SHC4 | ENSG00000185634 |
| | UNC5C | ENSG00000182168 |

| | | |
|---|---|---|
| | GABRA2 | ENSG00000151834 |
| | GRIA4 | ENSG00000152578 |
| | GABRA4 | ENSG00000109158 |
| | GABRG2 | ENSG00000113327 |
| | CHRNA4 | ENSG00000101204 |
| | GLRA3 | ENSG00000145451 |
| | CHRM2 | ENSG00000181072 |
| | GAD1 | ENSG00000128683 |
| | UNC13C | ENSG00000137766 |
| | CDH10 | ENSG00000040731 |
| | SYNDIG1 | ENSG00000101463 |
| | PPFIBP2 | ENSG00000166387 |
| | LRRTM1 | ENSG00000162951 |
| | PCLO | ENSG00000186472 |
| | DSCAM | ENSG00000171587 |
| | KCNJ3 | ENSG00000162989 |
| | IL1RAPL1 | ENSG00000169306 |
| | SLITRK2 | ENSG00000185985 |
| | SLITRK1 | ENSG00000178235 |
| | SLC8A3 | ENSG00000100678 |
| | ASIC2 | ENSG00000108684 |
| | TENM2 | ENSG00000145934 |
| | SLC4A10 | ENSG00000144290 |
| | SH3GL3 | ENSG00000140600 |
| | SH3GL2 | ENSG00000107295 |
| | CADPS2 | ENSG00000081803 |
| | TANC1 | ENSG00000115183 |
| | FAM107A | ENSG00000168309 |
| | NRN1 | ENSG00000124785 |
| | PDLIM4 | ENSG00000131435 |

| | | |
|---|---|---|
| | SNCB | ENSG00000074317 |
| | SNCG | ENSG00000173267 |
| | CACNA1A | ENSG00000141837 |
| | SYNPO | ENSG00000171992 |
| | SH2D5 | ENSG00000189410 |
| | KCNA3 | ENSG00000177272 |
| | KCNN2 | ENSG00000080709 |
| | HTR5A | ENSG00000157219 |
| | SYNC | ENSG00000162520 |
| Synapses - vesicles | SV2C | ENSG00000122012 |
| | SLC17A6 | ENSG00000091664 |
| | SLC6A17 | ENSG00000197106 |
| | WASF2 | ENSG00000158195 |
| Synapses - transmission | SYT7 | ENSG00000011347 |
| | SYT2 | ENSG00000143858 |
| | FRMPD4 | ENSG00000169933 |
| Golgi - glycosylation | GALNT9 | ENSG00000182870 |
| | GALNTL6 | ENSG00000174473 |
| | GALNT13 | ENSG00000144278 |
| | BGN | ENSG00000182492 |
| | DCN | ENSG00000011465 |
| | MAN1C1 | ENSG00000117643 |
| | XYLT2 | ENSG00000015532 |
| | UGT8 | ENSG00000174607 |
| | PLOD3 | ENSG00000106397 |
| | MGAT4C | ENSG00000182050 |
| | MGAT3 | ENSG00000128268 |
| | CHST9 | ENSG00000154080 |
| | SDC4 | ENSG00000124145 |
| | B4GALT1 | ENSG00000086062 |

| | | |
|---|---|---|
| | BCAN | ENSG00000132692 |
| | FUT1 | ENSG00000174951 |
| | OGN | ENSG00000106809 |
| | HSPG2 | ENSG00000142798 |
| | A4GALT | ENSG00000128274 |
| | CLEC18B | ENSG00000140839 |
| Golgi - organization/dynamics | FYCO1 | ENSG00000163820 |
| | GORASP1/2 | ENSG00000114745 |
| | GOLGA2/GM130 | ENSG00000167110 |
| Golgi - transport | COPZ2 | ENSG00000005243 |
| | RAB13 | ENSG00000143545 |
| | RAB34 | ENSG00000109113 |
| | RAB11FIP5 | ENSG00000135631 |
| | RAB7B | ENSG00000276600 |
| | PLIN3 | ENSG00000105355 |
| | SORL1 | ENSG00000137642 |

**Table S1D.** List of selected differentially expressed genes identified by bioinformatic analyses of RNA sequencing. Overview of selected differentially expressed genes identified through RNA sequencing and bioinformatic analyses, including involved compartment/pathway, gene name and ensemble ID.

| Target Gene | Ensemble ID | Forward Primer | Reverse Primer |
|---|---|---|---|
| CKMT1A | ENSG00000223572 | GATTCTGCCGAGGCCTCAAA | CCAGTGCCCAGGTTAGATGG |
| SCARA3 | ENSG00000168077 | CCGGAACCTCTCCATGATCG | GCCCTTTCACGCCCATATCT |
| FYCO1 | ENSG00000163820 | GGCCCCAGAAAGTTTCGGTT | CCTTGTTCTGTGGGCACTCT |
| GORASP | ENSG00000114745 | AGGTCTGGGAAGGGGATATGA | TCCACAATCTCTCGATGCCG |
| GOLGA2 | ENSG00000167110 | CTAGCGGTAGCCCTGGACTC | CCCTGCTTTTGGTGGCATTC |
| SORL1 | ENSG00000137642 | GGATACGGACTGCCAGGATG | AGCAATCACGCAGACCATCA |
| SYNDIG1 | ENSG00000101463 | GCAGCCTTCTACTTGTCCCA | CACATAGACGCCAGTCCCAA |
| GRIA4 | ENSG00000152578 | AGGTGAATGTGGACCCAAGG | CAAGCCGCCAACCAGAATGT |

| | | | |
|---|---|---|---|
| SYT7 | ENSG00000011347 | ACGAAGGGGACCATGTACCG | CCGCAGAGGACGATAGTGAC |
| GAPDH | ENSG00000111640 | CTCTCTGCTCCTCCTGTTCGAC | TGAGCGATGTGGCTCGGCT |

**Table S1E.** List of primers used for qPCR. Overview of differentially expressed target genes validated by qPCR, including gene name and ensemble ID, as well as forward and reverse primer design.

### Supplementary Experimental Procedures

#### hiPSC Generation and Cell Culture

The hiPSC fAD cell lines were derived from a patient carrying the fAD-linked A79V *PSEN1* mutation (3) and its gene corrected isogenic control A79V GC (4), a patient carrying the fAD-linked L150P *PSEN1* mutation (1) and its gene corrected isogenic control L150P GC (2), a healthy control (K3P53, (5)) and a CRISPR/Cas9 gene edited knock-in *APP* Swedish fAD line (BioSweden, (6)). All fAD lines have previously been published (Table S1A). The sAD cell lines have not been previously published but have been characterized and assessed for pluripotency markers and genomic integrity (Figure S2). The hiPSCs were cultured in Essential 8 (E8) media (A1517001, Thermo Fisher Scientific) on Matrigel-coated plates (TH Gayer, 7643022), and media was replaced every day. The cells were passaged every third day for approximately two weeks, before neural induction was initiated.

#### Neural Induction and Differentiation

Once hiPSC reached approximately 90% confluence, neural differentiation was performed according to a modified dual SMAD protocol (7). Neural induction was initiated by changing the media into neural induction media containing 50% DMEM/F12 (Thermo Fisher Scientific, 11330057), 50% advanced neurobasal medium (Thermo Fisher Scientific, 21103049), 1% N2 (Thermo Fisher Scientific, 17502048), 1% B27 without retinoic acid (Thermo Fisher Scientific, 1258010), 1% Glutamax (Thermo Fisher Scientific, 35050061), 1% non-essential amino acid

(NEAA, Thermo Fisher Scientific, 11140-050), 0,1% Pen/Strep (Sigma, P0781-100ML), supplemented with the inhibitors 10 µM SB431542 (SMAD inhibitor, SMS-gruppen, S1067) and 0,1 µM LDN193189 (Noggin analog, Sigma-Aldrich, SML0559). The cells were maintained in induction media for 12 days with daily media change. On day 12, a uniform neuroepithelial sheet appeared, and the neural progenitor cells (NPCs) were passaged with Accutase (Thermo Fisher Scientific, A1110501) into neural expansion media containing growth factors 10 ng/ml FGF2 (ProSpec, CYT-557) and 10 ng/ml EGF (ProSpec, CYT-217) instead of the inhibitors. NPCs were expanded and banked. Following expansion, NPCs were plated onto Poly-L-Ornithine (PLO, Sigma-Aldrich, P4957)/laminin (Sigma-Aldrich, L2020-1mg) coated dishes with a seeding density of 50 000 cells/cm$^2$, and terminal neural differentiation was performed in neural maturation media, supplemented with 50 µM db-cAMP (Sigma Aldrich, D0627-100mg), 200 µM Ascorbic acid (Sigma Aldrich, A4403-100MG), 20 ng/ml BDNF (ProSpec, CYT-207) and 10 ng/ml GDNF (ProSpec, CYT-305). The maturation process was carried out for five weeks for MitoTracker™ and Golgi ICC analysis and seven weeks for assessment of Aβ secretion and Tau phosphorylation as well as MitoTracker™, Golgi and synaptic evaluation, with partial media change every third day, before the neurons were fixed or harvested for further analyses.

**Mesoscale Assessment of Aβ Peptide Secretion**

Secretion of Amyloid beta (Aβ) peptides; Aβ38, Aβ40 and Aβ42, was assessed using the V-PLEX Plus Aβ Peptide Panel 1 Kit (MSD, 6E10), and the MESO QUICKPLEX SQ 120 Imager (MSD), connected to the DISCOVERY WORKBENCH 4.0 software, according to the manufacture's protocol. The data was normalized to the RNA concentration of the individual samples.

**Immunocytochemistry and Confocal Microscopy**

For immunocytochemistry (ICC), neurons were plated and cultured on PLO/laminin-coated, double acid-treated coverslips for five weeks for MitoTracker$^{TM}$ and Golgi assessment and seven weeks for neural verification, MitoTracker$^{TM}$, Golgi and synaptic evaluation, fixed in 4% Paraformaldehyde (PFA) for 20 minutes at room temperature. Following, neurons were washed 3x5 minutes in PBS (Sigma, D8537). Neurons were then permeabilized with 0.2% Triton-X-100 solution for 20 minutes at room temperature, then blocked with 3% Bovine Serum Albumin (BSA, Sigma) for 30 minutes at room temperature. Neurons were incubated with primary antibodies (Table S1B), diluted in 3% BSA, overnight at 4°C, washed with PBS 3x5 minutes at room temperature, and incubated with secondary antibodies (Table S1B) for 1 hour in the dark at room temperature. Another 3x5 minutes wash with PBS was performed, followed by DNA labelling with DAPI, diluted in PBS, for 7 minutes dark at room temperature. The neurons were then washed 3x5 minutes in PBS, before coverslips were mounted onto slides in DAKO fluorescence mounting solution and analyzed with the Leica confocal TCS SPE Microsystem implementing LAS x software. Obtained images were then processed in Fiji ImageJ 2.0.0-rc-65/1.51s.

**Neurite Outgrowth Analysis**

After five days of maturation, neurons were fixed in 4% PFA and ICC was performed with Beta-III-Tubulin (Tuj1, Millipore, MAB1637). Neurite length was measured and analyzed using Neurite Tracer in Fiji ImageJ (8).

**MitoTracker™ Assay**

Neurons were plated on coverslips, cultured for seven weeks, then incubated with 50 nM MitoTracker™ Red CMXRos (Invitrogen/Molecular Probes, M7512) in DMEM/F12 for 15 min at 37°C. Following, the neurons were fixed in 4% PFA, permeabilized in 0.2% Triton X-100 in PBS, incubated with DAPI for DNA labelling, washed and then mounted onto slides in mounting media. Neurons were imaged by confocal microscopy and analyzed in Fiji ImageJ. For detailed description see section: Immunocytochemistry and Confocal Microscopy.

**Transmission Electron Microscopy**

Neurons were cultured on coverslips, and after seven weeks of maturation fixed with 3% Gluteraldehyde (Merck, 1042390250) in 0.1 M Na-phosphate buffer with pH 7.2 at 4°C for 1 hour. The neurons were then embedded in 4% agar and cut into 1-2 $mm^3$ blocks under a stereomicroscope, then washed with 0.1 Na-phosphate buffer, followed by post-fixation in 1% osmium tetroxide (EMS) in 0.1% Na-phosphate buffer for 1 hour at room temperature. Washing with MilliQ water was performed, followed by a stepwise dehydration in ethanol with increasing concentration. Propylene oxide (Merck) was used as an intermediate allowing for infiltration with Epon (TAAB, T031). The following day, neurons were embedded in Epon, and cured at 60°C for 48 hours. Semi thin sections (2 µm) were cut on an ultramicrotome with glass knives (Leica Ultracut, Leica Microsystems, Wetzlar, Germany), then contrasted with 1% Toluidine blue (Millipore, 1159300025) in 1% Borax (LabChem, LC117101). Ultra-thin sections (50-70 nm) were cut on the microtome with a diamond knife (Jumdi, 2 mm), and the sections were collected onto grids, and then stained with 2% uranyl acetate (Polyscience) and lead citrate (Reynolds 1963). Imaging and analysis were performed "blinded" using a Philips CM100 transmission electron microscope with a Morada digital camera equipment and iTEM software system (Olympus).

Morphometry was used for quantitative evaluation of mitochondria, and 5 grids from each cell line were used for the analysis. 15 spots were randomly chosen at low magnification and positions were stored according to their X and Y coordinates using FEI/Philips CompuStage.

Images of each spot were captured using high magnification (19,000X), resulting in 75 images per cell line, and a total of 300 images. Three mitochondria categories were established based on the qualitative evaluation: normal mitochondria with well-defined cristae throughout the organelle, cristaeless mitochondria where these structures were lacking, and an additional category with mitochondria not fitting either of the two former categories (undefined). Mitochondria morphology and size were evaluated, and Fiji ImageJ was used for analysis by application of a grid to each image. Intersections were counted for each mitochondria category, as well as the cytoplasm and the number of mitochondria to calculate the relative mitochondria/cytoplasm ratio and the relative individual mitochondria area.

**Western Blot**

Neurons were cultured for seven weeks, washed with PBS, then lysed in M-PER mammalian protein extraction reagent (Thermo Fisher Scientific, 78501) with cOmplete™ Protease Inhibitor (Roche, 11873580001) and PhosSTOP™ phosphatase inhibitor (Roche, 04906845001). Protein content was determined using the Bradford Assay (Sigma, B6916). Separation of 10 µg protein was performed by NuPAGE™ Novex™ 4-12% Bis-Tris mini gel (Thermo Fisher Scientific, NP0335BOX), and immunoblotted with primary antibodies (Table S1C) overnight at 4°C, followed by secondary antibody incubation (Table S1C). The immunoblots were developed using Odyssey® Fc Imaging System (LI-COR) and analyzed using ImageStudio version 5.2.5 and Excel. Expression levels of target proteins were normalized against housekeeping proteins (GAPDH or β-actin).

**Amyloid Beta Treatment**

To evaluate the implication of Amyloid beta (Aβ) on the neurons, both the isogenic controls (A79V GC and L150P GC) and healthy control (K3P53) were treated with 2.5 µM Aβ peptide

1-42 (Sigma, PP69-0.25MG), incubated at 37°C for 24 hours, then fixed and harvested for ICC and TEM.

**RNA Extraction for bulk RNA Sequencing**

RNA was extracted from L150P and L150P GC as well as A79V and A79V GC neurons after seven weeks of neural maturation, using the RNeasy® Plus Mini Kit (Qiagen, 74134), according to the manufacturer's protocol, and the RNA quality was assessed using an Agilent 2100 Bioanalyzer system with RNA 6000 nano chip and reagents. Library preparation and sequencing (DNBseq, 2x100 nt, stranded paired-end) was done by an external provider for the fAD and isogenic control neurons (Beijing Genomics Institute, BGI).

**Analysis of bulk RNA Sequencing data**

The computational analysis of the RNA sequencing data is described in detail in a separate bioinformatics publication (Corsi, Gadekar et al, in prep.). Briefly, reads were pre-processed with Cutadapt v1.18 (9) to trim low quality 3' ends (min. Phred = 30), remove residual adapters, and filter out resulting reads shorter than 20 nt (pair_filter = "any"). Sequences matching to rRNAs in SILVA v119.1 (10) were removed with BBDuk v38.22 (http://jgi.doe.gov/data-and-tools/bb-tools/). Genes were quantified from the reads pairs aligned with STAR v.2.6.1d (11) on the human genome (hg38) using featureCounts (subread v.1.6.3, min. overlap 90 nt, min. fraction overlapping nt 90%, both aligned reads pairs non-chimeric) (12) provided with an extended set of annotations obtained by merging the GENCODE (13) and the FANTOM-CAT gene models (14). Differential expression analysis was carried out using DESeq2 v1.22.2 (15) and genes with Benjamini-Hochberg (16) adjusted Wald test P-value $< 0.05$, absolute log2 fold change $> 1$, and mean of normalized counts $> 10$ were considered significant. The workflow includes RseQC v.4.0.0 (17) for the confirmation of the library strandness and the verification of the editing status at on-target and potential off-target sites with CRISPRroots v.1.2 (18). Potential off-targets identified by CRISPRroots as high-risk were validate by sequencing, and no off-target edit was found (Corsi, Gadekar et al., in prep.).

**qPCR Validation**

RNA from L150P and L150P GC, A79V and A79V GC as well as BioSweden and K3P53 hiPSC derived neurons after seven weeks of neural maturation was extracted using the RNeasy® Plus Mini Kit (Qiagen, 74134). cDNA was synthesized from 1 μg of total RNA, according to Promega ImProm-II™ Reverse Transcription System (Promega, A3800). RNA was mixed with 1:3 oligoT: random primer (0.5 μg/μl), heated at 70°C for 5 minutes, then immediately put on ice for 5 minutes. Reverse transcription mix containing ImProm II buffer, dNTP mix (10mM), Nuclease-free water, RNasin Ribonuclease inhibitor (40u/μl), $MgCl_2$ (2.5 mM) and ImProm II was added, following 5 minutes incubation at room temperature, then 1 hour at 42°C. The enzyme was inactivated at 70°C for 15 minutes, and appropriate dilutions (1:5) of cDNA were prepared for further qPCR.

For qPCR analysis, nuclease-free water, Quantifast SYBRGreen 2x (Qiagen), forward- and reverse primers (10 μM, Table S1E) were mixed and added to each well of a qPCR plate, with a total of 8 μl per reaction and 2 μl of diluted cDNA was added. The qPCR was performed in triplicates, and a non-template control without cDNA was included for each target gene. The analysis was run on a LightCycler® 480 real-time PCR system (Roche, Switzerland) with a total of 40 cycles, and the data was collected and processed using the Design & Analysis software 2.6.0. The data were normalized to *GAPDH* gene.

**Proteomic Assessment**

For proteomic assessment, cell pellets from L150P and L150P GC as well as A79V and A79V GC neurons were collected using ice-cold PBS and a cell scraper. Following, proteomic assessment was performed by mass spectrometry, according to a previously published protocol (19).

***N*- and *O*-glycan profiling by mass spectrometry**

From L150P and L150P GC, A79V and A79V GC as well as BioSweden, K3P53 and Aβ-treated K3P53 hiPSC derived neurons, total cell lysate protein *N*- and *O*-glycans were released and analyzed by C18 nanoflow liquid chromatography (LC) coupled to mass spectrometry (MS) as described previously, with minor adaptations (20). Cell pellets were resuspended at ~$2x10^4$ cells/µL in lysis buffer (50 mM Tris HCl, 100 mM NaCl and 1x cOmplete™ protease inhibitor (EDTA-free)) and loaded on a preconditioned PVDF membrane. *N*-glycans were released using 2 U PNGase F in 30 µL water, eluted and dried. Sialic acids were derivatized by ethyl esterification (α2,6-linked sialic acids) and subsequent ammonia amidation (α2,3-linked sialic acids) (21) and purified by hydrophilic interaction liquid chromatography (HILIC) solid phase extraction (SPE) (22). Next, 50 µL 2-aminobenzamide (2-AB) reagent (500 mM 2-AB, 116 mM 2-methylpyridine borane complex (PB) in 45:45:10 metanol:water:acetic acid) was added and the samples were incubated 2.5 h at 50 °C. The glycans were purified by HILIC SPE and eluted in 50 µL water. Ten microliters of the eluates were diluted in 10 µL water for MS analysis. *O*-glycans were released from the same samples on the PVDF membrane using 20% hydroxylamine and 20% 1,8-diazabicyclo(5.4.0)undec-7-ene (DBU) for 1 h at 37 °C, and enriched by hydrazide beads, 2-AB labelled as described above, and purified by HILIC and porous graphitized carbon (PGC) SPE (20). Samples were resolved in 20 µL water for MS analysis. For both the *N*- and *O*-glycan preparations of each sample, 2 µL was injected for nanoLC-MS/MS analysis, using a single analytical column setup. The analytical column was prepared using a PicoFrit Emitter (New Objectives, 75 µm inner diameter), packed with Reprosil-Pure-AQ C18 phase (Dr. Maisch, 1.9 µm particle size, 22-25 cm column length). The emitter was interfaced to an Orbitrap Fusion Lumos mass spectrometer (Thermo Fisher Scientific) via a nanoSpray Flex ion source. Samples were eluted in an 1 h method with a gradient from 3% to 32% of solvent B in 35 min, from 32% to 100% B in the next 10 min and 100% B for the last 15 min at 200 nL/min (solvent A: 0.1% formic acid in water; solvent B: 0.1% formic acid in 80% ACN). A precursor MS scan (m/z 200-1700, positive polarity) was acquired in the Orbitrap at a nominal resolution of 120,000, followed by Orbitrap higher-energy C-trap dissociation (HCD)-MS/MS at a nominal resolution of 50,000 of the 10 most abundant precursors in the MS spectrum (charge states 1 to 4). A minimum MS signal threshold of 30,000 was used to trigger data-dependent fragmentation events. HCD was performed with an energy of 27% ± 5%, applying a 20 s dynamic exclusion window. Data analysis and structural annotation were performed as described before (20) using the Minora Feature Detector node in Thermo Proteome Discoverer 2.2.0.388 (Thermo Fisher Scientific lnc.), GlycoWorkbench 2.1 (build 146), (23) Skyline 21.1.0.146 (ProteoWizard) (24) and the Thermo Xcalibur qual browser 3.0.63. MS/MS spectra were manually assigned for each MS1 feature in at least one sample. *N*- and *O*- glycans were relatively quantified separately, by total area normalization. Derived traits were calculated based on specific glycosylation features, including for *N*-glycans the glycan type (pauci mannose, oligomannose, complex or hybrid) and complex-type fucosylation (no or core and/or antenna), sialylation (no or α2,3- and/or α2,6-linked), bisection, LacdiNAc, galactosylation and antenarity. For the *O*-glycans, the relative abundance of the *O*-glycan types was determined (*O*-GlacNAc, *O*-GalNAc, *O*-fucose, *O*-glycose or *O*-mannose), and specifically for the *O*-GalNAc glycans the relative abundance of the different cores (Tn, 1,

2 or 3) as well as the level of sialylation per core type. All values per cell type were represented as averages and standard deviations over three technical replicates.

**Airyscan super-resolution microscopy and Analysis**

Neuro 2A (N2A) cells were transiently transfected with either *APP* Swedish- or wild type h*APP*, using Lipofectamine 2000 (Thermo Fisher Scientific, 11668030) following a similar protocol as described previously (25). Briefly, after incubation, diluted DNA and Lipofectamine were combined and incubated for 20 minutes at room temperature to establish DNA-Lipofectamine complexes. The complex (500 µl) was then added to the N2A cells, which were further incubated for 24-72 hours at 37°C. Media was replaced after 6 hours. Following transfection, the N2A cells were fixed and immunocytochemical labelling for GM130 and γ-adaptin (Table S1B) was performed. The N2A cells were mounted with Prolong (Molecular Probes, cat. no. MAN0010261) for Airyscan super-resolution microscopy. Airyscan super-resolution microscopy was performed as described previously (26). Airyscan images were obtained on Zeiss LSM 880 equipped with 32 array detectors for acquisition of super-resolution images. For image acquisition, 405, 488, and 633 nm lasers were used. The illumination parameters like intensities, digital and analogue gain of the detectors, sampling of the images, emission window for each fluorescent channel and their corresponding pinhole sizes were maintained constant across acquisition paradigm. The raw images acquired using Airyscan mode were processed using Zeiss ZEN 3.4 (Blue) software to generate final super-resolution images. The reconstruction parameters were also kept constant throughout the samples. The morphological and biophysical traits of cis-/trans-Golgi were quantified on super-resolution images through MetaMorph software (Molecular Devices).

**5xFAD Transgenic Mice**

To further validate our findings, male 5xFAD transgenic mice (N=5) and male wild-type mice (N=4) were sacrificed, and brains were dissected. The cortex and hippocampus were fixed in 3% Gluteraldehyde and embedded in Epon for TEM evaluation. For detailed description of TEM preparation and imaging, see the section Transmission Electron Microscopy.

**Statistical Analysis and Data Availability**

Statistical analyses were performed using GraphPad Prism (Version 9.2.0) with default options, and statistical significance was determined using a Student's *t* test, multiple *t* test or two-way Anova with correction for multiple comparison. Data is presented as mean ± standard error of the mean (SEM) for all experiments with statistical significance *p < 0.05, **p < 0.01, ***p < 0.001 and ****p < 0.0001.

22. Selman MHJ, Hemayatkar M, Deelder AM, Wuhrer M. Cotton HILIC SPE microtips for microscale purification and enrichment of glycans and glycopeptides. Anal Chem

[Internet]. 2011 Apr 1 [cited 2022 Feb 26];83(7):2492–9. Available from: https://pubmed.ncbi.nlm.nih.gov/21366235/

23. Damerell D, Ceroni A, Maass K, Ranzinger R, Dell A, Haslam SM. The GlycanBuilder and GlycoWorkbench glycoinformatics tools: updates and new developments. Biol Chem [Internet]. 2012 Nov [cited 2022 Feb 26];393(11):1357–62. Available from: https://pubmed.ncbi.nlm.nih.gov/23109548/

24. Adams KJ, Pratt B, Bose N, Dubois LG, St. John-Williams L, Perrott KM, et al. Skyline for Small Molecules: A Unifying Software Package for Quantitative Metabolomics. J Proteome Res [Internet]. 2020 Apr 3 [cited 2022 Feb 26];19(4):1447–58. Available from: https://pubmed.ncbi.nlm.nih.gov/31984744/

25. Kedia S, Ramakrishna P, Netrakanti PR, Jose M, Sibarita JB, Nadkarni S, et al. Real-time nanoscale organization of amyloid precursor protein. Nanoscale [Internet]. 2020 Apr 17 [cited 2022 Jun 30];12(15):8200–15. Available from: https://pubs.rsc.org/en/content/articlehtml/2020/nr/d0nr00052c

26. Kedia S, Ramakrishna P, Netrakanti PR, Singh N, Sisodia SS, Jose M, et al. Alteration in synaptic nanoscale organization dictates amyloidogenic processing in Alzheimer's disease. iScience [Internet]. 2021 Jan 22 [cited 2022 Jun 30];24(1):101924. Available from: http://www.cell.com/article/S2589004220311214/fulltext

### Supplementary Information - Source Data

| Assessment | Cell line | Mean ± SEM (7 weeks) |
|---|---|---|
| Western blot (Ser199/Ser202) | L150P | 3.179 ± 1.6 |
| | L150P GC | 1.108 ± 0.1253 |
| | A79V | 1.423 ± 0.6141 |
| | A79V GC | 0.0301 ± 0.01727 |
| Western blot (T181) | L150P | 3.462 ± 0.3366** |
| | L150P GC | 0.9135 ± 0.1931 |
| | A79V | 0.036 ± 0.01069 |
| | A79V GC | 0.021 ± 0.01002 |
| Western blot (S422) | L150P | 2.718 ± 0.4762* |
| | L150P GC | 0.3592 ± 0.1359 |
| | A79V | 2.28 ± 0.9796 |
| | A79V GC | 2.2 ± 1.113 |
| Western blot (Ser396) | L150P | 1.135 ± 0.203 |
| | L150P GC | 1.098 ± 0.3337 |
| | A79V | 2.98 ± 0.6145 |
| | A79V GC | 0.4967 ± 0.0865 |
| Aβ40 (Mesoscale) | L150P | 34.42 ± 1.334**** |
| | L150P GC | 54.34 ± 0.9868 |
| | A79V | 36.03 ± 1.164**** |
| | A79V GC | 15.13 ± 0.6698 |
| | BioSWEDEN | 92.99 ± 1.855**** |
| | K3P53 | 202.5 ± 0.2531 |
| Aβ42 (Mesoscale) | L150P | 5.132 ± 0.1064**** |
| | L150P GC | 7.210 ± 0.02053 |

| | | |
|---|---|---|
| | A79V | 6.946 ± 0.1206**** |
| | A79V GC | 2.767 ± 0.03049 |
| | BioSWEDEN | 13.30 ± 0.4567**** |
| | K3P53 | 20.38 ± 0.1889 |
| Aβ42/Aβ40 ratio | L150P | 0.1497 ± 0.002891** |
| | L150P GC | 0.1329 ± 0.002744 |
| | A79V | 0.1942 ± 0.009176 |
| | A79V GC | 0.1845 ± 0.007334 |
| | BioSWEDEN | 0.1428 ± 0.002615**** |
| | K3P53 | 0.1006 ± 0.0008222 |
| Neurite outgrowth (Length) | L150P | 799.28 ± 126.53 |
| | L150P GC | 937.77 ± 135.48 |
| | A79V | 176.74 ± 26.7364 |
| | A79V GC | 196.82 ± 13.7152 |
| Synaptic density (SY38 puncta) | L150P | 23.47 ± 0.9735*** |
| | L150P GC | 29.47 ± 1.14 |
| | A79V | 14.96 ± 0.7703** |
| | A79V GC | 19.43 ± 1.293 |
| Total mitochondria (TEM) | L150P | 0.0365 ± 0.002 |
| | L150P GC | 0.034 ± 0.0032 |
| | A79V | 0.0382 ± 0.002 |
| | A79V GC | 0.0308 ± 0.0051 |
| Normal mitochondria (TEM) | L150P | 0.01539 ± 0.0014 |
| | L150P GC | 0.01998 ± 0.002 |
| | A79V | 0.01846 ± 0.0023 |
| | A79V GC | 0.0192 ± 0.0037 |
| Cristaeless mitochondria (TEM) | L150P | 0.01077 ± 0.0011* |
| | L150P GC | 0.00667 ± 0.0012 |

| | | |
|---|---|---|
| | A79V | 0.01814 ± 0.0024* |
| | A79V GC | 0.00856 ± 0.0023 |
| Mitochondria size (TEM) | L150P | 1.1921 ± 0.04695 |
| | L150P GC | 1.2459 ± 0.0518 |
| | A79V | 1.4373 ± 0.03689 |
| | A79V GC | 1.4199 ± 0.0466 |

**Table S2A.** Statistical analyses of observed cellular phenotypes in seven-week hiPSC derived neurons. Statistical analyses of observed cellular phenotypes in seven-week neurons, including western blot for Tau analysis, mesoscale for Aβ secretion, neurite outgrowth assessment, synaptic density assessment and mitochondria TEM morphometry analysis. Results are displayed as mean ± standard error of the mean (SEM) from three replicates.

| **Assessment** | **Cell line** | **Mean ± SEM (5 weeks)** | **Mean ± SEM (7 weeks)** |
|---|---|---|---|
| MitoTracker™ (Mean area of distribution) | L150P | 1472 ± 482.5 | 800.9 ± 233.3** |
| | L150P GC | 2603 ± 1004 | 2249 ± 309.1 |
| | A79V | 1358 ± 110.9 | 808.4 ± 172.7* |
| | A79V GC | 1285 ± 99.88 | 2671 ± 666.0 |
| | BioSWEDEN | - | 745.7 ± 55.06* |
| | K3P53 | - | 920.1 ± 54.97 |
| Cis-Golgi surface area ICC (GM130) | L150P | 256.5 ± 98.01 | 115 ± 12.00** |
| | L150P GC | 150.4 ± 55.29 | 28 ± 7.9 |
| | A79V | 187.2 ± 31.73* | 140.31 ± 13.65 |
| | A79V GC | 35.83 ± 4.550 | 108.9 ± 24.1 |
| | BioSWEDEN | - | 272.0 ± 29.92** |
| | K3P53 | - | 167.1 ± 13.88 |
| | L150P GC (Aβ) | - | 348.8 ± 51.89**** |
| | A79V GC (Aβ) | - | 152.0 ± 28.74 |
| Trans-Golgi surface area | L150P | 61.63 ± 7.473 | 36 ± 4.43* |

| | | | |
|---|---|---|---|
| ICC (TGN38) | L150P GC | 35.97 ± 16.80 | 22 ± 0.50 |
| | A79V | 240.0 ± 40.49* | 49.3 ± 8.2436* |
| | A79V GC | 35.97 ± 16.80 | 25.56 ± 5.009 |
| | BioSWEDEN | - | 371.0 ± 34.25**** |
| | K3P53 | - | 80.84 ± 5.848 |
| | L150P GC (Aβ) | - | 234.3 ± 24.02**** |
| | A79V GC (Aβ) | - | 118.0 ± 11.85**** |

**Table S2B.** Statistical analyses of mitochondria and Golgi alteration in five- and seven-week hiPSC derived neurons. Statistical analyses of mitochondria and Golgi alteration in five- and seven-week neurons. Results are displayed as mean ± standard error of the mean (SEM) from three replicates.

| Gene | Cell Line | RNA-seq analysis Log2 Fold Change | RNA-seq analysis Adjusted p-value | Proteomics analysis Abundance ratio | Proteomics analysis Adjusted p-value |
|---|---|---|---|---|---|
| SCARA3 | L150P | -2.43 | 9.89e-39 | 0,447 | 4.84e-07 |
| | A79V | -2.19 | 6.93e-20 | 0,658 | 8.18e-06 |
| ACSF2 | L150P | -1.14 | 4.22e-24 | 0,79 | 0.0002 |
| | A79V | -1.93 | 4.18e-51 | 0,669 | 7.27e-06 |
| NME4 | L150P | -1.18 | 5.34e-13 | 0,828 | 0.16 |
| | A79V | -1.15 | 1.62e-18 | 0,788 | 0.02 |
| ACSS1 | L150P | -2.80 | 5.39e-24 | 1,311 | 0.0079 |
| | A79V | -2.52 | 1.49e-15 | 0,562 | 2.28e-05 |
| ACSL6 | L150P | 1.85 | 2.15e-25 | 1,645 | 1.19e-06 |
| | A79V | 1.797 | 5.23e-15 | 1,275 | 0.0001 |
| DGAT2 | L150P | 1.17 | 2.66e-09 | - | - |
| | A79V | 1.29 | 4.75e-11 | - | - |
| ME1 | L150P | -1.02 | 8.53e-25 | 0,531 | 6.00e-06 |
| | A79V | -1.81 | 6.01e-09 | 0,633 | 4.21e-05 |
| ACADL | L150P | -2.12 | 8.01e-33 | 1,172 | 0.009 |
| | A79V | 1.65 | 7.30e-08 | 0,744 | 0.0001 |

| | | | | | |
|---|---|---|---|---|---|
| ACOT11 | L150P | 1.02 | 0.015 | - | - |
| | A79V | -1.55 | 0.0002 | - | - |
| CKMT1B | L150P | 2.004 | 1.03e-29 | 1,697 | 5.13e-07 |
| | A79V | 1.03 | 3.32e-09 | 1,122 | 0.002 |
| CKMT1A | L150P | 1.59 | 2.02e-18 | 1,697 | 5.13e-07 |
| | A79V | 1.01 | 1.127e-08 | 1,122 | 0.002 |
| SNAP25 | L150P | 0.95 | 1.49e-46 | 1,661 | 2.94e-05 |
| | A79V | 0.72 | 0.001 | 1,154 | 0.01 |
| GRM7 | L150P | 1.90 | 1.85e-120 | 2,31 | 4.74e-07 |
| | A79V | 0.20 | 0.35 | 1,278 | 0.0001 |
| GRIA1 | L150P | -1.03 | 4.06e-110 | 0,803 | 7.72e-06 |
| | A79V | 1.35 | 3.70e-55 | 1,272 | 4.70e-06 |
| SYAP1 | L150P | -0.24 | 0.03 | 1,08 | 0.02 |
| | A79V | -0.21 | 0.03 | 0,992 | 0.97 |
| SYP | L150P | 0.61 | 5.54e-06 | 1,693 | 2.24e-05 |
| | A79V | 1.24 | 1.41e-09 | 1,247 | 0.001 |
| GRIN1 | L150P | 1.10 | 4.70e-06 | - | - |
| | A79V | 0.82 | 0.0002 | - | - |
| CACNG5 | L150P | 2.92 | 2.28e-150 | - | - |
| | A79V | 3.24 | 2.70e-42 | - | - |
| CHRNA6 | L150P | 1.13 | 0.0001 | - | - |
| | A79V | 1.96 | 2.71e-25 | - | - |
| SHC4 | L150P | 1.36 | 1.24e-32 | - | - |
| | A79V | 1.85 | 1.37e-24 | - | - |
| UNC5C | L150P | -1.08 | 1.17e-29 | 1,052 | 1 |
| | A79V | -1.77 | 6.70e-20 | 0,675 | 0.0001 |
| GABRA2 | L150P | 1.35 | 9.87e-37 | 1,776 | 3.16e-06 |
| | A79V | 1.58 | 2.12e-19 | 1,135 | 0.01 |
| GRIA4 | L150P | 1.05 | 1.12e-18 | 1,091 | 0.06 |

| | | | | | |
|---|---|---|---|---|---|
| | A79V | 1.58 | 8.92e-19 | 1,1 | 0.02 |
| GABRA4 | L150P | 2.00 | 7.07e-36 | - | - |
| | A79V | 1.39 | 9.93e-12 | - | - |
| GABRG2 | L150P | 1.74 | 1.92e-76 | 2,049 | 4.37e-07 |
| | A79V | 1.46 | 1.76e-11 | 1,339 | 2.66e-05 |
| CHRNA4 | L150P | 1.16 | 2.71e-08 | - | - |
| | A79V | 1.39 | 6.49e-10 | - | - |
| GLRA3 | L150P | 1.36 | 6.98e-16 | - | - |
| | A79V | 1.28 | 1.42e-09 | - | - |
| CHRM2 | L150P | 3.46 | 1.25e-69 | - | - |
| | A79V | 1.17 | 2.83e-09 | - | - |
| GAD1 | L150P | 1.69 | 1.74e-83 | 0,786 | 0.001 |
| | A79V | 1.38 | 2.13e-31 | 1,315 | 0.0007 |
| UNC13C | L150P | 1.54 | 2.59e-25 | - | - |
| | A79V | 1.26 | 2.55e-13 | - | - |
| CDH10 | L150P | 1.29 | 2.86e-48 | 0,762 | 0.05 |
| | A79V | 1.51 | 1.70e-21 | 1,055 | 0.90 |
| SYNDIG1 | L150P | -1.28 | 7.60e-22 | - | - |
| | A79V | -1.47 | 1.76e-19 | - | - |
| PPFIBP2 | L150P | 1.30 | 4.43e-12 | - | - |
| | A79V | -1.79 | 1.11e-16 | - | - |
| LRRTM1 | L150P | 1.25 | 2.40e-28 | - | - |
| | A79V | 1.22 | 5.49e-14 | - | - |
| PCLO | L150P | 1.06 | 2.62e-13 | - | - |
| | A79V | 1.49 | 3.08e-19 | - | - |
| DSCAM | L150P | 1.55 | 9.75e-45 | - | - |
| | A79V | 1.02 | 3.99e-18 | - | - |
| KCNJ3 | L150P | 1.80 | 5.60e-43 | - | - |
| | A79V | 1.69 | 7.01e-16 | - | - |

| | | | | | |
|---|---|---|---|---|---|
| IL1RAPL | L150P | 1.21 | 7.39e-23 | - | - |
| 1 | A79V | 1.08 | 4.73e-12 | - | - |
| SLITRK2 | L150P | 3.08 | 1.15e-221 | 1,567 | 3.48e-06 |
| | A79V | 1.03 | 1.61e-10 | 1,169 | 0.004 |
| SLITRK1 | L150P | 1.45 | 1.44e-87 | 1,476 | 2.26e-06 |
| | A79V | 1.13 | 3.74e-09 | 1,215 | 0.0002 |
| SLC8A3 | L150P | 1.56 | 4.63e-85 | - | - |
| | A79V | 1.16 | 2.54e-10 | - | - |
| ASIC2 | L150P | 1.31 | 6.21e-09 | - | - |
| | A79V | 1.36 | 3.92e-10 | - | - |
| TENM2 | L150P | -1.24 | 1.09e-64 | 1,012 | 0.96 |
| | A79V | 1.20 | 8.59e-10 | 1,495 | 1.28e-06 |
| SLC4A10 | L150P | 1.10 | 2.04e-14 | 1,019 | 0.33 |
| | A79V | 1.02 | 2.19e-09 | 1,121 | 0.0008 |
| SH3GL3 | L150P | 1.08 | 2.14e-29 | 1,844 | 8.94e-07 |
| | A79V | 1.18 | 5.07e-09 | 1,117 | 0.02 |
| SH3GL2 | L150P | 1.83 | 3.33e-178 | 2,123 | 4.849e-07 |
| | A79V | 1.16 | 5.79e-07 | 1,262 | 0.0003 |
| CADPS2 | L150P | 2.78 | 3.26e-232 | 1,204 | 0.004 |
| | A79V | -1.21 | 7.06e-09 | 2,264 | 1.15e-06 |
| TANC1 | L150P | -1.78 | 1.13e-105 | - | - |
| | A79V | -1.009 | 2.03e-08 | - | - |
| FAM107A | L150P | 1.12 | 1.43e-24 | - | - |
| | A79V | 1.50 | 2.56e-08 | - | - |
| NRN1 | L150P | 2.92 | 2.05e-161 | 2,211 | 5.503e-06 |
| | A79V | 1.05 | 2.62e-08 | 1,421 | 0.001 |
| PDLIM4 | L150P | -1.14 | 1.85e-05 | 0,338 | 1.56e-06 |
| | A79V | -1.56 | 3.09e-08 | 0,456 | 1.82e-05 |
| SNCB | L150P | 1.23 | 3.30e-08 | 1,713 | 4.14e-05 |

| | | | | | |
|---|---|---|---|---|---|
| | A79V | 1.33 | 6.77e-08 | 1,211 | 0.03 |
| SNCG | L150P | 2.41 | 1.64e-05 | 1,617 | 7.65e-06 |
| | A79V | 1.01 | 0.002 | 1,243 | 0.001 |
| CACNA1A | L150P | 1.57 | 1.40e-37 | - | - |
| | A79V | 1.37 | 7.91e-08 | - | - |
| SYNPO | L150P | -3.92 | 1.85e-185 | 0,427 | 5.13e-07 |
| | A79V | -1.58 | 1.26e-06 | 0,712 | 7.76e-05 |
| SH2D5 | L150P | 1.28 | 1.32e-07 | - | - |
| | A79V | 1.19 | 2.10e-05 | - | - |
| KCNA3 | L150P | 1.15 | 1.68e-14 | 1,085 | 0.19 |
| | A79V | 1.02 | 2.90e-05 | 1,324 | 0.006 |
| KCNN2 | L150P | 3.63 | 1.11e-29 | - | - |
| | A79V | 1.007 | 8.07e-05 | - | - |
| HTR5A | L150P | 1.39 | 0.0005 | - | - |
| | A79V | 1.15 | 0.0003 | - | - |
| SYNC | L150P | -2.51 | 2.22e-249 | 0,496 | 2.00e-06 |
| | A79V | -1.40 | 0.01 | 0,774 | 0.0006 |
| SV2C | L150P | 3.09 | 0.0 | - | - |
| | A79V | -1.36 | 2.16e-19 | - | - |
| SLC17A6 | L150P | 2.85 | 3.17e-146 | 2,896 | 6.21e-07 |
| | A79V | 1.44 | 1.64e-14 | 1,392 | 0.0006 |
| SLC6A17 | L150P | 1.46 | 7.29e-22 | 1,559 | 5.98e-07 |
| | A79V | 1.47 | 4.13e-13 | 1,333 | 5.61e-06 |
| WASF2 | L150P | -1.26 | 6.29e-142 | 0,462 | 5.44e-07 |
| | A79V | -1.24 | 2.84e-12 | 0,67 | 2.28e-05 |
| SYT7 | L150P | 1.08 | 9.13e-17 | 2,257 | 1.98e-06 |
| | A79V | 1.06 | 1.98e-08 | 1,059 | 0.40 |
| SYT2 | L150P | 3.46 | 1.93e-46 | 1,315 | 0.01 |
| | A79V | 2.26 | 3.05e-26 | 1,422 | 0.002 |

| | | | | | |
|---|---|---|---|---|---|
| FRMPD4 | L150P | 1.26 | 8.34e-21 | - | - |
| | A79V | 1.20 | 1.94e-06 | - | - |
| GALNT9 | L150P | 1.52 | 8.91e-07 | - | - |
| | A79V | 1.46 | 2.36e-11 | - | - |
| GALNTL | L150P | 1.57 | 4.95e-40 | - | - |
| 6 | A79V | 1.04 | 0.0002 | - | - |
| GALNT13 | L150P | 1.96 | 3.21e-58 | - | - |
| | A79V | 1.13 | 1.49e-15 | - | - |
| BGN | L150P | -2.82 | 4.96e-27 | 1,154 | 0.17 |
| | A79V | -2.60 | 3.85e-121 | 0,408 | 4.16e-05 |
| DCN | L150P | 3.23 | 5.23e-65 | 1,143 | 0.37 |
| | A79V | -3.19 | 4.62e-120 | 0,362 | 9.68e-05 |
| MAN1C1 | L150P | -1.88 | 1.14e-73 | 0,528 | 2.03e-07 |
| | A79V | -1.45 | 1.00e-32 | 0,59 | 3.30e-07 |
| XYLT2 | L150P | -1.03 | -4.72 | 1,092 | 0.43 |
| | A79V | -1.29 | 4.51e-16 | 0,952 | 0.99 |
| UGT8 | L150P | 2.14 | 9.46e-46 | - | - |
| | A79V | -2.69 | 6.89e-16 | - | - |
| PLOD3 | L150P | -1.51 | 1.46e-12 | 0,54 | 4.07e-07 |
| | A79V | -1.12 | 9.02e-13 | 0,592 | 5.77e-07 |
| MGAT4C | L150P | 1.54 | 2.75e-23 | - | - |
| | A79V | 1.65 | 1.14e-12 | - | - |
| MGAT3 | L150P | 0.58 | 0.003 | - | - |
| | A79V | 0.24 | 0.04 | - | - |
| CHST9 | L150P | -1.43 | 7.22e-20 | - | - |
| | A79V | 1.07 | 6.92e-12 | - | - |
| SDC4 | L150P | -1.81 | 3.12e-77 | 1,286 | 0.07 |
| | A79V | -1.97 | 2.11e-10 | 1,251 | 0.13 |
| B4GALT1 | L150P | -1.25 | 8.33e-42 | - | - |

| | | | | | |
|---|---|---|---|---|---|
| | A79V | -1.70 | 4.87e-09 | - | - |
| BCAN | L150P | 1.34 | 1.56e-12 | 1,94 | 1.18e-05 |
| | A79V | 1.81 | 4.69e-08 | 1,512 | 0.0001 |
| FUT1 | L150P | 1.37 | 1.004e-10 | - | - |
| | A79V | 1.40 | 3.873e-07 | - | - |
| OGN | L150P | -1.22 | 1.58e-05 | 1,065 | 0.52 |
| | A79V | -2.13 | 5.88e-07 | 0,422 | 1.91e-05 |
| HSPG2 | L150P | -1.68 | 1.46e-10 | 0,939 | 0.06 |
| | A79V | -1.44 | 3.25e-06 | 0,87 | 0.002 |
| A4GALT | L150P | -3.34 | 1.09e-12 | - | - |
| | A79V | -2.58 | 9.15e-06 | - | - |
| CLEC18B | L150P | -2.75 | 1.74e-32 | 0,791 | 0.01 |
| | A79V | -1.57 | 0.009 | 0,931 | 0.27 |
| FYCO1 | L150P | -1.30 | 7.85e-51 | 0,795 | 0.0001 |
| | A79V | -1.38 | 1.62e-18 | 0,754 | 3.24e-05 |
| GORASP | L150P | -0.37 | 0.001 | 0,978 | 1 |
| 1/2 | A79V | -0.28 | 1.23e-07 | 1,002 | 1 |
| GOLGA2/ | L150P | -0.18 | 0.14 | 1,021 | 0.27 |
| GM130 | A79V | -0.80 | 2.42e-56 | 0,82 | 1.15e-05 |
| COPZ2 | L150P | -2.30 | 1.05e-41 | 0,866 | 0.02 |
| | A79V | -3.31 | 3.67e-78 | 1,375 | 0.50 |
| RAB13 | L150P | -1.01 | 8.52e-17 | 0,591 | 5.21e-05 |
| | A79V | -1.58 | 1.66e-15 | 0,95 | 0.23 |
| RAB34 | L150P | -1.19 | 1.05e-18 | 0,61 | 4.53e-06 |
| | A79V | -1.79 | 3.94e-15 | 0,519 | 1.13e-06 |
| RAB11FI | L150P | -1.01 | 3.34e-09 | 0,961 | 0.87 |
| P5 | A79V | -1.04 | 3.87e-14 | 1,178 | 0.01 |
| RAB7B | L150P | -1.69 | 1.66e-19 | - | - |
| | A79V | -1.63 | 2.93e-13 | - | - |

| | | | | | |
|---|---|---|---|---|---|
| PLIN3 | L150P | -1.30 | 1.17e-12 | 0,536 | 1.22e-06 |
| | A79V | -1.13 | 1.82e-08 | 0,411 | 5.34e-07 |
| SORL1 | L150P | 0.48 | 8.98e-13 | 1,042 | 0.69 |
| | A79V | 1.26 | 8.50e-19 | 1,05 | 0.195 |

**Table S2C.** RNA sequencing (RNA-seq) and proteomics statistics. RNA sequencing (RNA-seq) and proteomics statistical data. RNA-seq data is displayed as log2 fold change and adjusted p-value, and proteomic data is demonstrated by abundance ration and adjusted p-value. – indicates no measured expression.
